# Compositionality of social gaze in the prefrontal-amygdala circuits

**DOI:** 10.1101/2025.07.28.667161

**Authors:** Guangyao Qi, Olga Dal Monte, Siqi Fan, Steve W. C. Chang

## Abstract

Social gaze underpins primate communication, yet the neural principles enabling its flexibility remain unknown. Each social gaze can be deconstructed into three primitives: gaze content, social state, and gaze duration. To reduce dimensionality and facilitate generalization, the brain needs to represent these primitives in an abstract format. Here we show that social gaze is governed by a compositional code built from these primitives in the brain. In male and female macaques (neural recordings in two males) engaged in real-life social gaze interaction, behavior analyses revealed that partner responses were determined by how primitives were combined, rather than by their independent sums, providing evidence for behavioral compositionality. The basolateral amygdala and the anterior cingulate gyrus represented content and state in an abstract format and orthogonally to one another, whereas the dorsomedial prefrontal and orbitofrontal cortices exhibited limited generalization. Linear mixed-selective neurons encoding both content and state in the basolateral amygdala and the anterior cingulate gyrus, but not in the other two areas, facilitated the abstraction underlying generalization. Moreover, distinct channels routed content and state information across prefrontal–amygdala circuits to minimize interference, which was mediated by linear mixed selectivity neurons. These findings identify a neural grammar for social gaze, revealing compositional computations as a principle of flexible social communication.

## Main

In humans and nonhuman primates, social behaviors are powerfully guided by gaze dynamics of the interacting individuals ^1,2^. In social gaze interaction, functionally unique components, or primitives, underlie each social gaze event, where different combinations of these gaze primitives produce drastically different functional meanings. Every social gaze event can be described with respect to multiple primitives, including the target of the social gaze (‘content’), the level of social engagement in which the gaze takes place as a proxy to an internal state guiding these social gaze behaviors ^3–5^ (‘state’), and the length of that gaze fixation (‘duration’). These primitives can be functionally separable ^6–8^ and, importantly, may underlie the flexibility found in social gaze interaction. For example, a quick look at the partner’s face during high engagement may serve as a cue for turn-taking, while the same gaze during low engagement might be interpreted as a mere monitoring behavior ^9^. Thus, the compositional nature of social gaze primitives may contribute to the complexity and richness of social gaze functions.

Compositional theories suggest that cognitive processes are built from shared components that can be flexibly recombined to generate a wide range of behaviors ^10–14^. However, it remains elusive if and how the brain computes social gaze primitives to support potential compositional processing. More specifically, it remains unknown whether structured relationships in firing patterns, such as representational neural geometry, exist among these primitives that could support compositional processing.

Importantly, compositional processes benefit from abstraction and orthogonal representations of primitives, as such representations can help reduce interference and enhance robustness to noise ^11,15,16^. How might the brain represent social gaze primitives in an abstract format that reduces dimensionality and facilitates generalization? To realize flexible control of social gaze interaction, the relevant neural populations not only need to distinctively represent social gaze information, but also structure the firing patterns of the population of neurons across different primitives through abstraction ^17^, with the ultimate goal of communicating the relevant information across neural networks.

Here we investigated the compositionality of social gaze by examining the neural representations of three specific social gaze primitives - gaze content, social state, and gaze duration - in the primate prefrontal-amygdala circuits during real-life social gaze interactions taking place between pairs of macaques ^6,18,19^. We applied representational geometry analysis to test the generalization of social gaze primitives, tested the single-neuron basis of generalization, and examined the directional flow of the content and state information in the prefrontal-amygdala networks. Neural populations in the basolateral amygdala and the anterior cingulate gyrus orthogonally represented content and state in an abstract, generalizable, format and maintained their separability, whereas neurons in the dorsomedial prefrontal cortex and the orbitofrontal cortex exhibited limited generalization. At the single-neuron level, linear mixed selectivity cells facilitated the abstraction underlying the generalization. Finally, we found distinct routing of content and state information across the four brain regions. Together, our results reveal novel insights into how the brain represents social gaze primitives to support compositional computations for guiding social gaze interaction.

## Results

### Combinations of social gaze primitives determine interactive social gaze behavior

We hypothesized that the brain represents social gaze in a compositional manner. To test this hypothesis, we first decomposed social gaze into three specific gaze primitives: content (i.e., target of the gaze), state (i.e., social engagement level at the time of the gaze), and duration (i.e., duration of the gaze) (Fig. 1A; Methods). We then examined how neural populations in four distinct brain regions in the primate prefrontal-amygdala circuits – the basolateral amygdala (BLA), anterior cingulate gyrus (ACCg), orbitofrontal cortex (OFC), and dorsomedial prefrontal cortex (dmPFC) – represented these primitives during social gaze interaction (Fig. 1B). In the experimental setting (Fig. S1), pairs of macaques (M1 with neural recording, and M2 serving as partner monkeys) freely interacted with gaze without any imposed task constraints.

**Figure 1.**
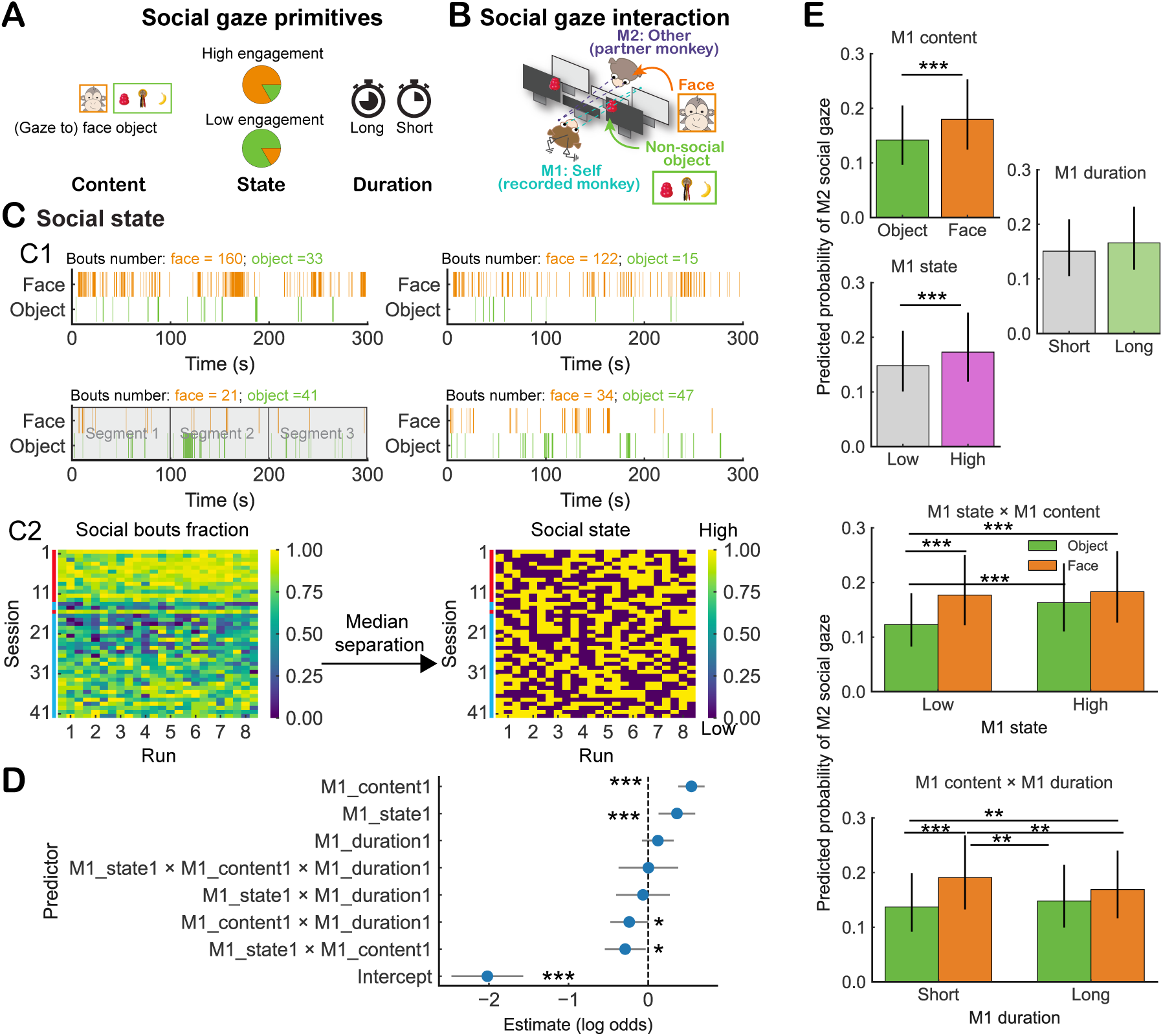
Social gaze primitives combinatorially guide real-life social gaze interaction. (**A**) Social gaze primitives encompass gaze content (identifying the subject of the gaze), social state (indicating the level of social engagement), and gaze duration (measuring how long the gaze persists). (**B**) Social gaze interaction paradigm. Pairs of monkeys sat face-to-face while both their eye positions were tracked. The recorded monkey (M1) freely gazed at the partner monkey (M2) or the object. (**C**) Social state. C1 upper: two example 5-min runs showing high social engagement with an elevated rate of gaze to the partner’s face over the object. C1 lower: two example runs showing low social engagement with an elevated rate of gaze to the object over the partner’s face. C1 lower left: an example of how each run was divided into multiple segments. C2: (Left) Normalized social gaze (face) bouts fraction across runs and sessions. (Right) Social states across runs and sessions based on median separation. Each run was equally divided into 100-sec segments, and the normalized social bouts were calculated for each segment. The vertical color bars on the left of each heatmap indicates the recorded subject (M1) identity across sessions (red marks: sessions 1–13 and 16 from Monkey L; cyan marks: sessions 14–15 and 17–42 from Monkey K). **(D)** Estimates of fixed effects from the generalized linear mixed-effects model (GLMM) predicting the occurrence of M2’s social gaze based on M1’s social gaze primitives and their compositions (Methods). Each point represents a fixed effect (log-odds estimate) with 95% confidence intervals. Asterisks indicate significance levels based on Type III Wald chi-square tests. The reference conditions for all predictors were defined as M1 state = Low, M1 content = Object, M1 duration = Short. **(E)** Predicted probabilities (back-transformed from the logit scale) of M2 social gaze as a function of M1 content, M1 state, M1 duration, and the compositions of these social gaze primitives. (Top three panels) Main effects for M1 content (Object vs. Face), M1 state (Low vs. High), and M1 duration (Short vs. Long). (Bottom two panels) Two-way interactions: M1 state × M1 content and M1 content × M1 duration. Predicted probabilities are shown by M1 state, separately by content (Object vs. Face). Vertical lines on the bars indicate 95% confidence intervals. Horizontal lines with asterisks denote significant pairwise differences. Significance was tested using pairwise Wald z-tests on marginal mean differences (Tukey-adjusted), which account for covariance; thus, significance may occur even when 95% confidence intervals overlap. * *P* < 0.05; ** *P* < 0.01; *** *P* < 0.001.

Over the course of a 40-min session in each day (8 runs, 5-min each), the frequency of gaze fixations on the face and the object robustly fluctuated over time (Fig. 1C1), likely signifying dynamic changes in social engagement, which we use to operationally define ‘social state’. Because social state is a covert, internal state that cannot be observed directly, we inferred it from overt behavior ^20^ by treating the moment-to-moment frequency of face- and object-directed gaze fixations as a behavioral proxy for the underlying level of social engagement. To quantify these fluctuations, we divided each 5-min run into three 100-sec segments resulting in a total of 24 segments per session. We then calculated normalized social gaze bouts as an index of social state for each segment (Fig. 1C2; Methods). We selected a 100-sec window because it offers the highest temporal resolution that still reliably captures social-state dynamics in the data, while ensuring each window contains at least one face-directed and one object-directed gaze bout for subsequent analyses. The social state index for each 100-sec window (segment) was defined as the proportion of face-directed gaze bouts out of the total number of face-and object-directed gaze bouts in that segment. The ‘high’ and ‘low’ social states were then categorized based on whether the social state index (Methods) was higher or lower than each session’s median social state index (Fig. 1C2). Notably, these classified states were not associated with pupil sizes (mean Fisher Z-transformed correlation = –0.0016; *P* = 0.97, one-sample *t*-test against zero; Fig. S2), suggesting that social engagement states do not simply index general arousal levels. Instead, these states may reflect a category of an internal state ^20^ that can vary independently from processes faithfully indexed by pupil dynamics ^21^. Similarly, the ‘high’ and ‘low’ gaze durations were categorized as whether the gaze duration index of each gaze event was higher or lower than each session’s median gaze duration index (Fig. S3A; Methods). Finally, the gaze content was determined by the target of each gaze event (Fig. S3B; face or object).

To examine behavioral correlates of compositionality in social gaze behavior, we asked how different combinations of gaze primitives of the recorded monkeys (M1) influenced the partner’s (M2) social gaze behaviors. Here we hypothesized that M1’s primitive components not only influence the partner’s social gaze behavior, but also distinct combinations of these primitives have different influences on them. If so, these behavioral results will support the productivity aspects of compositionality.

To test these hypotheses, we applied a generalized linear mixed-effects model (GLMM) to predict the probability of M2’s gaze to M1’s face (M2’s social gaze) based on different combinations of M1’s social gaze primitives (Methods). This model included fixed effects for M1 content, M1 state, and M1 duration, as well as all possible two-way and three-way interactions among these primitives. We observed significant main effects of M1 content (z = 6.47, *P* < 0.001, Wald z-test) and M1 state (z = 3.08, *P* = 0.002) on the probability of M2’s social gaze occurrence, while M1 duration alone exhibited a trend effect on this probability (z = 1.20, *P* = 0.23) (Fig. 1D). Crucially, in addition to the main effects, we observed significant interactions – there was a negative interaction between M1 state and M1 content (z = –2.21, *P* = 0.027), indicating that the combined effect of these two primitives was sublinear (i.e., smaller than the linear sum of their individual effects). Further, M1 content and M1 duration also showed a significant interaction (z = –1.98, *P* = 0.048), indicating that the influence of M1 content on M2’s social gaze probability was modulated by the temporal characteristics of M1’s fixation duration.

Predicted probabilities from the GLMM revealed that the partner monkeys were more likely to initiate social gaze (gaze to M1’s face) when M1 was in a high social state (*P* < 0.001, Tukey-adjusted) and when M1 looked at M2’s face (*P* < 0.001) (Fig. 1E, top panel). By contrast, the influence of M1 duration on M2’s gaze behavior was not reliable (*P* = 0.54). Interaction analyses revealed that the combination of high M1 state and M1 face content did not further amplify M2’s social gaze probability as expected under simple additivity (Fig. 1E, bottom panel). Specifically, when M1 was fixating on the object, M2 was more likely to look at M1 if M1 was in a high social state compared to a low social state (*P* < 0.001). However, when M1 was fixating on the face, M2’s likelihood of looking at M1 did not differ between high and low social states of M1 (*P* = 0.624). Moreover, compared to M1’s object-directed gaze, M1’s face-directed gaze generally encouraged M2 to look at M1 (Fig. 1E, top left), but this increase in M2’s social gaze was selective to when M1 was in a low state (*P* < 0.001 vs. *P* = 0.282 when M1 was in a high state; Fig. 1E, bottom left). Importantly, when the model was fit separately for each M1 monkey, the direction of the main effects was consistent across animals (content: β (log-odds) = 0.62 and 0.25; state: β = 0.36 and 0.35 for Monkeys K and L, respectively). Consistent with the pooled analysis, both the state × content and the content × duration interactions were negative in both monkeys (state × content: β = −0.01 and −0.49; content × duration: β = −0.28 and −0.22 for Monkeys K and L, respectively).

Crucially, the selection of gaze content, social state, and gaze duration as primitives is not based on conceptual considerations alone (Methods), but is supported by explicit behavioral model comparisons. To test whether these variables constitute behaviorally dissociable components, we evaluated whether each primitive contributed non-redundant explanatory information beyond the others. Using the full model (Fig. 1D) as the baseline, we then performed block-wise likelihood-ratio tests to assess primitive non-redundant (Fig. S4). For each primitive (gaze content, social state, gaze duration), we constructed a reduced model in which the main effect of that primitive and all associated interaction terms were jointly removed, while all remaining terms were retained. Each reduced model was compared directly to the full model using a likelihood-ratio test. Removing gaze content resulted in a large and highly significant degradation of model fit (Δ−2 log L = 58.36, df = 4, *P* = 6.4 × 10⁻¹²). Importantly, removing social state or gaze duration also significantly reduced model fit (state: Δ−2 log L = 16.08, df = 4, *P* = 0.0029; duration: Δ−2 log L = 16.28, df = 4, *P* = 0.0027; Fig. S4A). These effects exceeded the χ² critical value (α = 0.05), indicating that each primitive explains variance in partner gaze behavior that cannot be absorbed by the remaining variables. We further evaluated model support using an information-theoretic framework. ΔAIC values relative to the full model increased substantially when any primitive was removed (Fig. S4B), indicating reduced model support in all three cases. Together, likelihood-ratio tests and AIC-based comparisons demonstrate that gaze content, social state, and gaze duration are behaviorally dissociable components of social gaze.

In a complementary analysis, we also examined how M1’s social gaze primitives predicted M2’s social state itself. Notably, M1’s social state exhibited a significantly negative association with M2’s social state (z = –2.85, *P* = 0.004, Wald z-test) (Fig. S5) such that when M1 looked at M2 more frequently (i.e., high M1 social state), M2 looked at M1 less often (low M2 social state), and vice versa. This finding aligns well with the tendency of rhesus macaques to observe each other’s faces while avoiding excessive direct eye contact ^18,22,23^.

In a control analysis, we asked whether the observed gaze coordination dynamics could be explained by stable dominance–subordination relationships between monkeys. To address this possibility, we extended the original generalized linear mixed-effects model to include partner dominance (a dyad-level, binary variable reflecting stable social hierarchy) as an additional predictor of M2’s social gaze behavior (Methods). This analysis revealed no significant main effect of dominance, nor any significant interactions between dominance and M1’s gaze content, state, or duration (all *P* > 0.1; Fig. S6A). Importantly, the previously reported effects of gaze content, gaze state, and their interaction—including the sublinear interaction between state and content—remained robust and unchanged after accounting for dominance. As an additional robustness check, we repeated the analysis treating dominance as a random effect rather than a fixed effect. Under this alternative specification (Fig. S6B), again consistent with our main behavior results (Fig. 1D), the main effects of gaze content, state, interaction between content and state, and interaction between content and duration were preserved, further confirming that the primary behavioral findings were not driven by dominance relationships.

Taken together, these behavioral findings first demonstrate that social gaze behaviors are shaped by the compositional structure of social gaze primitives. Social state and gaze content play distinct roles in shaping the partner’s social gaze behavior. Specifically, partner’s social gaze behaviors were nonlinearly mapped to the combinations of social gaze primitives, supporting unique functional correlates with respect to how the social gaze primitives are combined.

### Neural coding of social gaze primitives in the prefrontal cortex and the amygdala supports the compositionality hypothesis

To test the neural encoding of each gaze primitive, we generated three hypothesized representational dissimilarity matrices (RDMs), in which each hypothesized RDM was orthogonal to the others (Fig. 2A). To examine the encoding of social gaze primitives in BLA, ACCg, dmPFC, and OFC in the RDM space (Fig. 2B), we generated the brain RDMs for each region by first averaging the firing rates of individual neurons within each gaze event condition (i.e., 8 combination conditions of gaze content, social state, and gaze duration; Methods) (Fig. 2C). All neural analyses were aligned to the onset of individual gaze events. For each event, neural activity was summarized as the average firing rate over the full duration of that gaze event (Methods). We then computed pairwise Euclidean distances between condition-level population vectors across the eight conditions (Methods). After generating brain RDMs, we used a generalized linear model (GLM) (Fig. 2D) to test the encoding of social gaze primitive RDMs.

**Figure 2.**
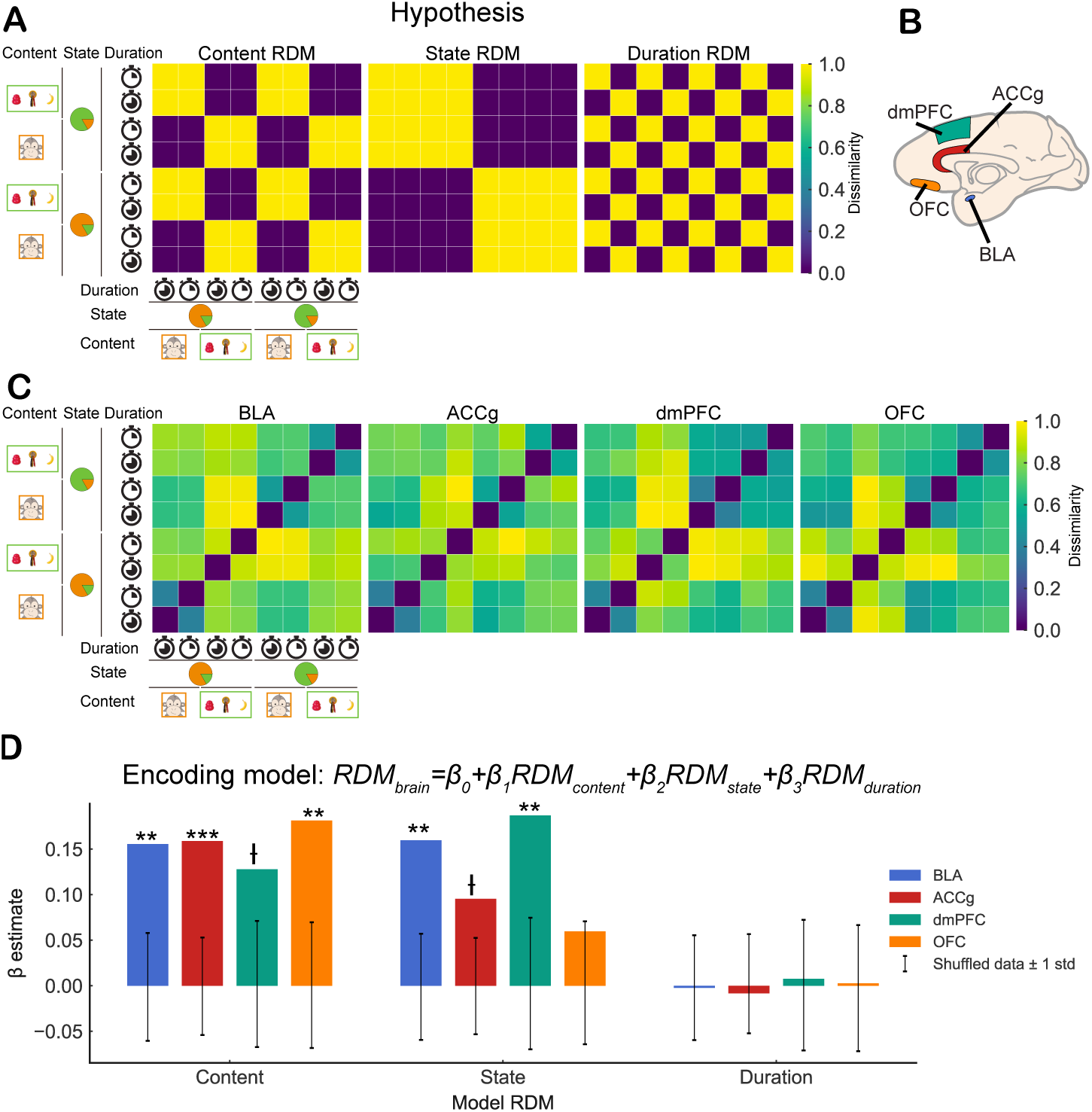
Encoding of social gaze primitives by the four neural populations in the prefrontal-amygdala circuits. (**A**) Three hypothesized representational dissimilarity matrices (RDMs) for testing the neural encoding of social gaze primitives. (**B**) An illustration of the four neural populations examined. ACCg, anterior cingulate gyrus; BLA, basolateral amygdala; dmPFC, dorsomedial prefrontal cortex; OFC, orbitofrontal cortex. (**C**) The neural RDM results from the recorded brain areas. (**D**) Encoding of the three social gaze primitives in the four neural populations based on the encoding GLM model (Methods). Ɨ *P* < 0.1; * *P* < 0.05; ** *P* < 0.01; *** *P* < 0.001 with permutation test.

According to the encoding model, the gaze content was widely encoded in all tested neural populations (Fig. 2D), with positive β estimates observed in both monkeys across regions (Monkey K: ACCg: 0.18; BLA: 0.17; dmPFC: 0.14; OFC: 0.12; Monkey L: ACCg: 0.12; BLA: 0.14; dmPFC: 0.11; OFC: 0.28). The social state information was encoded in BLA, ACCg, and dmPFC but the evidence of that was only weak in OFC (Fig. 2D), again showing consistent effect directions across monkeys (Monkey K: ACCg: 0.096; BLA: 0.18; dmPFC: 0.16; OFC: 0.073; Monkey L: ACCg: 0.086; BLA: 0.12; dmPFC: 0.21; OFC: 0.046). However, none of the four brain regions reliably represented gaze duration (Fig. 2D), even when models were fit separately for each monkey (Monkey K: ACCg: −0.011; BLA: −0.0058; dmPFC: 0.0031; OFC: −0.0041; Monkey L: ACCg: −0.0016; BLA: 0.0037; dmPFC: 0.012; OFC: 0.010), with duration β estimates clustering near zero (approximately −0.01 to 0.01), despite the duration had interactive behavioral effects with the content and state (Fig. 1D). We found similar results with more strict regression analyses (Fig. S7). We will therefore focus on the representation of state and content in the four neural populations. Motivated by this qualitative consistency in both behavioral structure and neural encoding across monkeys, we pooled neural data across subjects for the following neural analyses to maximize statistical power.

A neural ensemble can encode the social gaze primitives through a variety of representational formats. Here we focused on examining evidence in support of the compositional processing of social gaze by testing whether the representations of social gaze primitives are distinct from one another and if the primitives can support generalization across conditions ^11,14^. Specifically, we tested three key predictions derived from the compositionality hypothesis ^13,24^ (Fig. 3A, Fig. S8): (*i*) that neural representations of social state and gaze content are distinct; (*ii*) that state and content are encoded along orthogonal axes; and (*iii*) that these representations generalize across conditions, such that state information can be used across different contents and content information can be used across different states (Fig. 3D top panel). To test these predictions, we used a population decoding-based geometry analysis ^25^ of social states and gaze content (Fig. 3A).

**Figure 3.**
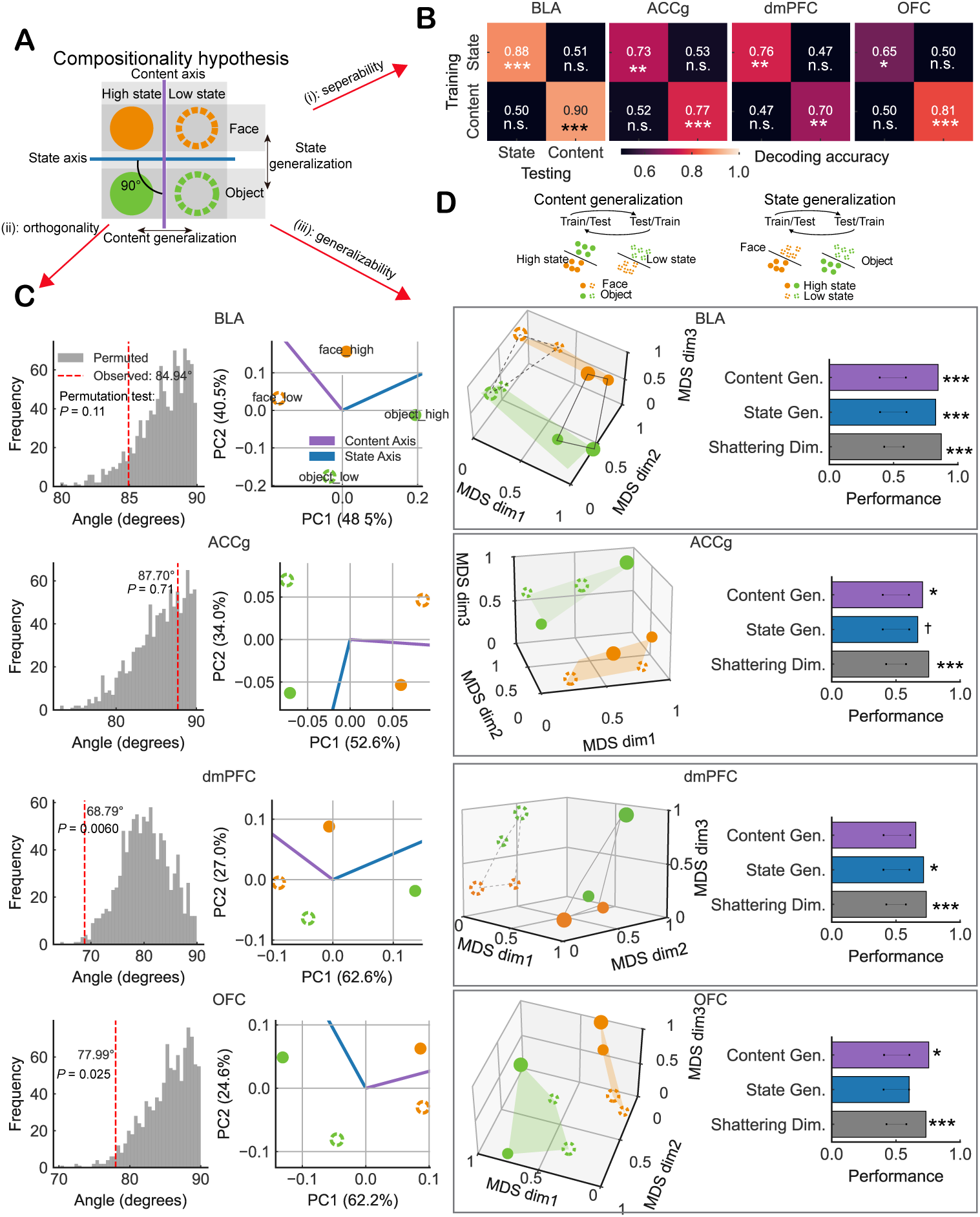
Tests of the three predictions from the compositionality hypothesis of social gaze in the prefrontal-amygdala circuits. **(A)** Schematic of a compositional encoding hypothesis, showing the three testable predictions. Neural population responses to four conditions (content/state: face/high, object/high, face/low, object/low) are distributed along orthogonal axes in neural state space. Color (orange = face, green = object) and fill (solid = high state, open = low state) distinguish the two dimensions. See Methods and Fig. S8 for more details on the three predictions from the hypothesis. (**B**) Cross-primitives decoding results. n.s., not significant; * *P* < 0.05; ** *P* < 0.01; *** *P* < 0.001 with permutation test. **(C)** Orthogonality tests between the content axis and state axis. Left: Angle between the gaze content coding axis and social state coding axis, estimated from condition differences (Methods). Right: Projection of mean firing rates during the gaze period onto a two-dimensional neural subspace spanned by the first two components from PCA. The dots indicate four different gaze events: face/high (gaze to face in high social state), face/low (gaze to face in low social state), object/high (gaze to object in high social state), and object/low (gaze to object in low social state). **(D)** The neural geometry of social gaze primitives in BLA, ACCg, dmPFC, and OFC. Top: Cross-conditions decoding for the generalization test. Top left, A linear SVM trained to decode face vs. object in one state condition (e.g., low state) successfully generalizes to the other state (e.g., high state), supporting the idea that content representations are reused across states (i.e., content generalization). Top right, A linear SVM trained to decode high vs. low state in one content condition (e.g., face) successfully generalizes to the other content (e.g., object), supporting the idea that state representations are reused across content (i.e., state generalization). Bottom left, Multidimensional scaling (MDS) analysis derived from the brain RDMs (Fig. 2C). Distances between points reflect a three-dimensional multidimensional scaling embedding of the neural representational dissimilarity matrix. Large and small circles indicate long and short gaze duration, respectively. Orange and green circles indicate face and object gaze content, respectively. Filled and hollow circles indicate high and low states, respectively. Content planes are color-filled, whereas state planes are outlined in solid and dashed lines. Bottom right, Cross-condition generalization performances of gaze content and social state, as well as the neural separability of any two conditions across four experimental conditions (content/state: face/high, face/low, object/high, and object/low). Content gen.: content generalization. State gen.: state generalization. Shattering dim.: shattering dimensionality. Error bar: shuffle data ± SD. n.s., not significant; Ɨ *P* < 0.1; * *P* < 0.05; ** *P* < 0.01; *** *P* < 0.001 with permutation test.

All four tested brain areas could reliably decode both gaze content and social state (Fig. 3B). Although we methodologically decomposed the primitives in an orthogonal manner (Fig. 2A), the neural representations for the content and state may still be representationally shared in these neural populations. To test if their neural representations are indeed not shared (prediction (*i*)), we applied a cross-primitives decoding analysis by training a linear gaze content decoder and testing it for social state, and vice versa (Fig. 3B; Methods). It was clear from this analysis that the decoder could not cross-decode different social gaze primitives. All the tested decoding accuracies were close to the chance level (Fig. 3B). The absence of cross-primitives decodability (test of prediction (*i*)) supports that gaze content and social state representations are not representationally shared.

Next, we tested the orthogonality between the neural representations of gaze content and social state (prediction (*ii*)) by performing a neural state space analysis ^26^ (Methods). For each brain region, condition-specific population vectors were derived from mean firing rates across four sub-conditions (face/object content × high/low social state). Two encoding axes – one for content and the other for state – were defined based on the average firing rate differences among relevant conditions. We then calculated the angle between these axes to quantify the degree of independence between the two representations. This orthogonality test directly quantifies the independence of the population codes for these two primitives, going beyond separability to test how distinct their encoding subspaces are. Only BLA and ACCg passed the orthogonality test (Fig. 3C; BLA: 84.94°, *P* = 0.11; ACCg: 87.70°, *P* = 0.71, permutation test). By contrast, dmPFC (68.79°, *P* = 0.006) and OFC (77.99°, *P* = 0.025) had less orthogonal representations. These results indicate that social state and gaze content representations are largely unshared in BLA and ACCg, in contrast to the more coupled representations observed in dmPFC and OFC.

Finally, to examine whether the neural representation of state or content is structured in an abstract format and can be generalized to new conditions (prediction (*iii*)) (e.g., a linear decoder for state in face-directed events can be used in unseen object-directed events), we performed a cross-condition generalization analysis of social gaze primitives (Fig. 3A). To test for content generalization, we trained a content decoder in one state (high or low social engagement) and tested the decoder in another untrained state (low or high, respectively). For state generalization, we trained a state decoder in one content (looking at face or object) and tested the decoder performance in another untrained content (object or face, respectively). The average testing decoding accuracy was used to measure generalization ability.

Multidimensional scaling (MDS) of the RDMs revealed that BLA represents social gaze primitives most abstractly compared to the other brain regions (Fig. 3D left panel). The explained variance of the 3D MDS embeddings was high across regions (BLA: r² = 0.86; ACCg: r² = 0.86; dmPFC: r² = 0.90; OFC: r² = 0.80), indicating that the plotted distances (Fig. 3D) closely reflect the underlying neural representational geometry. In these multidimensional scaling results, conditions that form more clearly separated and planar groupings suggest more abstract and lower-dimensional representations. BLA encoded both content and state in distinct low-dimensional subspaces (Fig. 3D left panel). By contrast, representations in ACCg, dmPFC, and OFC were less abstract, with less aligned subspaces for content or state (Fig. 3D left panel). The quantifications of the generalization results indicate that both BLA and ACCg could generalize gaze content and social state (i.e., high content and state generalization performance in Fig. 3D right panel), whereas dmPFC and OFC could only generalize social state and gaze content, respectively (i.e., high state-only or content-only generalization) (Fig. 3D right panel). Furthermore, the shattering dimensionality analysis, another way to measure the dimensionality of neural geometry ^25^, demonstrated that all four brain regions did not reduce their separability for each sub-condition (high state-face, low state-face, high state-object, low state-object) (Fig. 3D right panel). This analysis was not designed to test decoding of specific condition pairs, but to quantify the overall capacity of population activity to support flexible linear separations across multiple condition boundaries, following the concept of shattering dimensionality. Taken together, BLA and ACCg represented gaze content and social state in an abstract format, maintaining their separability, while dmPFC and OFC were only able to generalize either social state or gaze content, but not both.

### Linear mixed selectivity neurons facilitate the generalization of social gaze primitives

Is there a relationship between single-neuron encoding and the population-level generalization of content and state? Previous work proposed that linear mixed selectivity neurons, whose activity reflects weighted combinations of multiple independent variables, may arise through abstraction and support generalization across conditions ^27^. These neurons can preserve condition-relevant information in low-dimensional but in flexible formats, which makes them ideal for supporting transferable computations. Identifying such neurons in the prefrontal-amygdala populations that generalize both content and state could therefore reveal the single-neuron basis for compositional representations of social gaze. Because only BLA and ACCg generalized *both* social state and gaze content together, we hypothesized that neurons with linear mixed selectivity in these regions support content and state generalization. To determine single-neuron encoding schema, we combined a two-step modeling approach: a standard linear regression (FR ∼ content + state) to assess main effects, followed by a comparison with a non-linear model including an interaction term (FR ∼ content + state + content:state) to test for the interaction (Fig. 4A; Methods). We classified each neuron into having (*i*) pure content selectivity, (*ii*) pure state selectivity, (*iii*) linear mixed selectivity, or (*iv*) nonlinear mixed selectivity (Fig. 4B, one proportion Z test with 0.05, *P* < 0.05). To assess potential collinearity between gaze content and social state in the single-neuron regression model, we quantified the variance inflation factors for the two predictors (content and state). VIF values for both predictors were close to 1 and well below the conventional threshold for problematic multicollinearity (VIF = 5) (Fig. S9), supporting the validity of the regression-based classification of neuronal selectivity.

**Figure 4.**
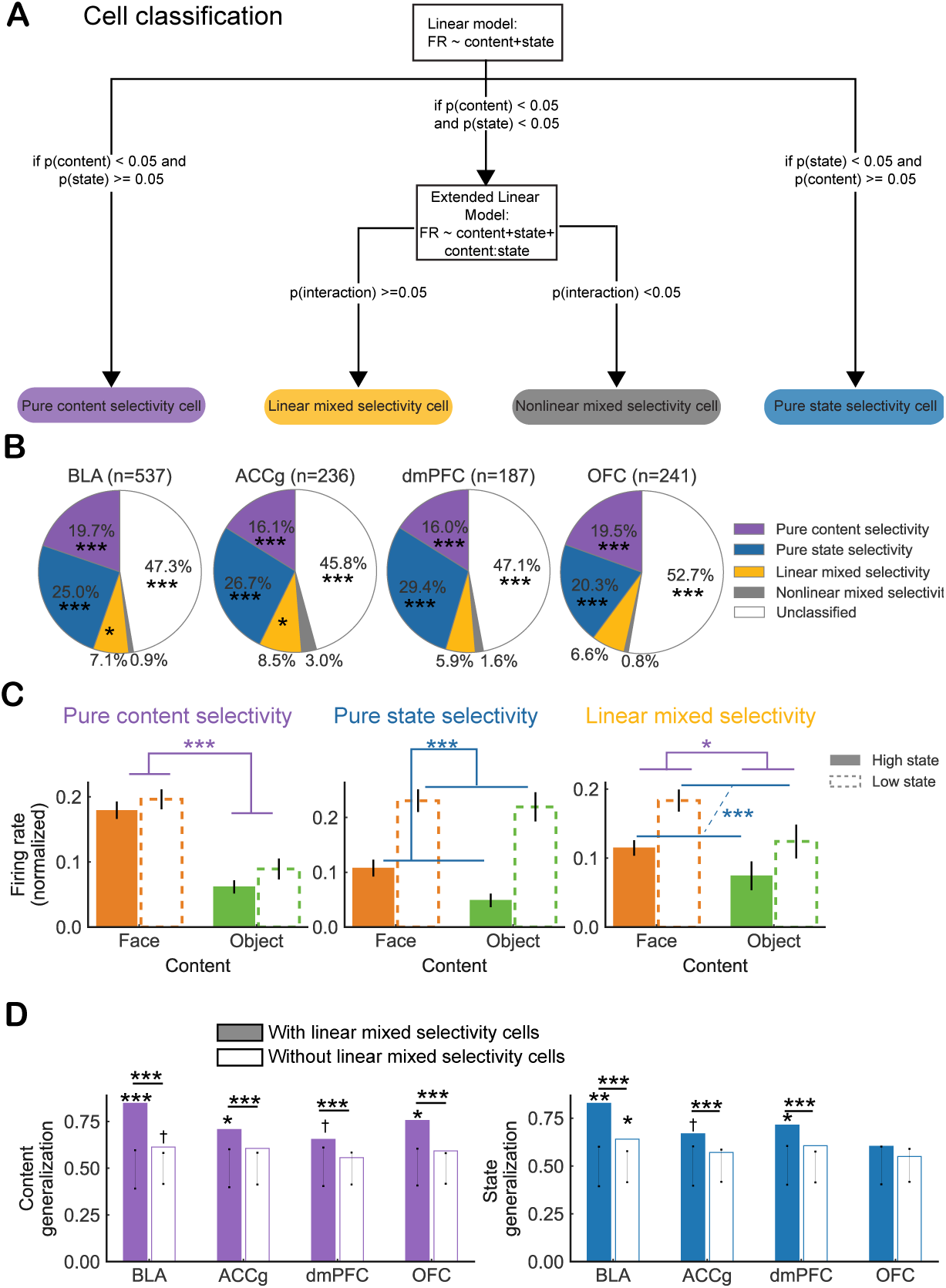
Contributions of linear mixed selectivity neurons to the generalization of social gaze primitives. **(A)** Diagram of the single-neuron classification procedure for identifying cells with pure selectivity, linear mixed selectivity, and nonlinear mixed selectivity. **(B)** Neuronal classifications from the four populations in the prefrontal-amygdala circuits. * *P* < 0.05; ** *P* < 0.01; *** *P* < 0.001 with one proportion Z test with 0.05. **(C)** Three example cells in BLA showing pure content selectivity (left), pure state selectivity (middle), and linear mixed selectivity (right). The peri-stimulus time histograms (PSTHs) of these same three BLA cells are shown in the upper panel of Fig. S10. Horizontal brackets indicate significant main effects of either gaze content (face vs. object, purple) or social state (high vs. low, blue) based on linear regressions in **(A)**. Error bar: mean ± SEM. * *P* < 0.05; *** *P* < 0.001. (**D**) Generalization performance of gaze content and social state (n-matched population), with or without linear mixed selectivity cells. Error bar: shuffle data ± SD. Permutation tests were used for each column. Wilcoxon rank-sum tests were used to compare the columns. Ɨ *P* < 0.1; * *P* < 0.05; ** *P* < 0.01; *** *P* < 0.001.

All four areas contained substantial proportions of pure content and pure state neurons (Fig. 4B-C; Fig. S10), consistent with their ability to decode both content and state (Fig. 3B). Consistent with our hypothesis, only BLA and ACCg showed a proportion of cells with linear mixed selectivity that was significantly above chance (Fig. 4B). To visualize the temporal emergence of population selectivity, we computed population PSTHs for neurons modulated by gaze content or social state, recoded by preference using a cross-validated procedure (Fig. S11). Across areas, state-related population responses showed a sustained separation between preferred and non-preferred conditions throughout the analyzed interval, whereas content-related responses differed before gaze onset and progressively converged following fixation, revealing distinct temporal profiles for the two gaze primitives (Fig. S11). To investigate whether linear mixed neurons specifically contribute to generalization, we compared the generalization ability for population when including versus excluding cells with linear mixed selectivity. Crucially, the cross-condition generalization performance for both gaze content and social state was significantly decreased in BLA and ACCg when removing the linear mixed selectivity cells (Fig. 4D), suggesting that linear mixed selectivity encoding in single cells facilitates the generalization of social gaze primitives.

To test whether population-level RSA and decoding results depended on neurons without detectable task-related modulation, we repeated all analyses after excluding neurons that showed no significant effects of gaze content or social state, or their interaction. The resulting encoding and decoding patterns were qualitatively unchanged across all four brain regions, indicating that the reported population-level effects are not driven by the inclusion of non-selective neurons (Figs. S12 and S13).

### Social state modulation is not explained by short-term visual stimulation history

To determine whether neural modulation attributed to social state could instead reflect differences in recent visual stimulation, we performed a control analysis explicitly modeling the effect of the immediately preceding gaze event (“history”) (Methods). For this analysis, history was defined as the category of the immediately preceding gaze event (face = 1, object = 0), state was defined as the current social state (high = 1, low = 0), and firing rate was defined as the mean neural response during the current gaze event (post-gaze window), using the same event-aligned firing-rate measure as in all other single-neuron analyses. For each neuron, we fit a linear model of the form FR ∼ state + history, retaining only neurons for which all four state × history combinations were observed.

Across all brain regions, a substantial fraction of neurons showed significant modulation by social state, whereas fewer neurons were significantly modulated by visual history (BLA: state = 192/537, 36%; history = 60/537, 11%; ACCg: state = 90/236, 38%; history = 31/236, 13%; P = 0.697; dmPFC: state = 68/187, 36%; history = 13/187, 7%;; OFC: state = 77/241, 32%; history = 18/241, 8%). Critically, neurons significant for state and those significant for history exhibited only chance-level overlap (BLA: overlap = 18/537, 3%; ACCg: overlap = 11/236, 5%; dmPFC: overlap = 4/187, 2%; OFC: overlap = 4/241, 2%). Hypergeometric tests futher revealed no enrichment of overlap between state- and history-significant neurons in any region (BLA: P = 0.871; ACCg: *P* = 0.697; dmPFC: *P* = 0.764; OFC: *P* = 0.884). Thus, neurons sensitive to social state and those sensitive to immediate visual history largely constituted distinct populations across all four areas.

Furthermore, among neurons significant for both predictors, the magnitudes of state-related and history-related effects were not correlated. Pearson correlations between the regression coefficients for state (β_state) and history (β_history) were non-significant in all regions (BLA: r = −0.180, *P* = 0.476; ACCg: r = −0.245, *P* = 0.467; dmPFC: r = 0.254, *P* = 0.746; OFC: r = 0.605, *P* = 0.395). This indicates that neurons exhibiting stronger state modulation did not systematically exhibit stronger history modulation.

Together, these results show that social state–related neural modulation cannot be explained by immediate visual stimulation history. Instead, social state and short-term visual history exert largely separable influences on neural activity, supporting the interpretation of social state as an independent contributor to single-neuron responses rather than a proxy for recent visual input.

### Distinct communication subspaces are used for different social gaze primitives in the prefrontal-amygdala circuits

The results so far inform how the prefrontal cortex and amygdala support the compositionality of social gaze at the levels of single neurons and populations. Ultimately, the flow of information across multiple brain areas is critical for guiding social behavior. Here we thus asked how the representations of social gaze primitives are communicated among the four areas to support compositional principles. We hypothesized that the prefrontal-amygdala networks route distinct primitive components via separate communication subspaces to minimize information interference and enable flexible composition.

To test this hypothesis, we conducted a canonical correlation analysis (CCA) to identify correlations within a ‘communication subspace’ among the four brain regions as an index of neural communication^28^. Importantly, for examining content information communication, we aligned the content information but misaligned the state information for the tested pairs of regions, and vice versa for examining state information communication. The zero-lagged CCA analysis indicated that the content communication was relatively dynamic and gaze event-dependent, in which the content communication only emerged during and after gazing at the target content (Fig. 5A upper panel). In some pairs of areas (dmPFC-BLA and OFC-ACCg), this communication was exclusively present while the monkeys gazed at the target content. By contrast, the state communication, between BLA and dmPFC and also between BLA and ACCg, was consistent over time, emerging before each gaze event and continuing after gazing at the target content (face or object) (Fig. 5A lower panel).

**Figure 5.**
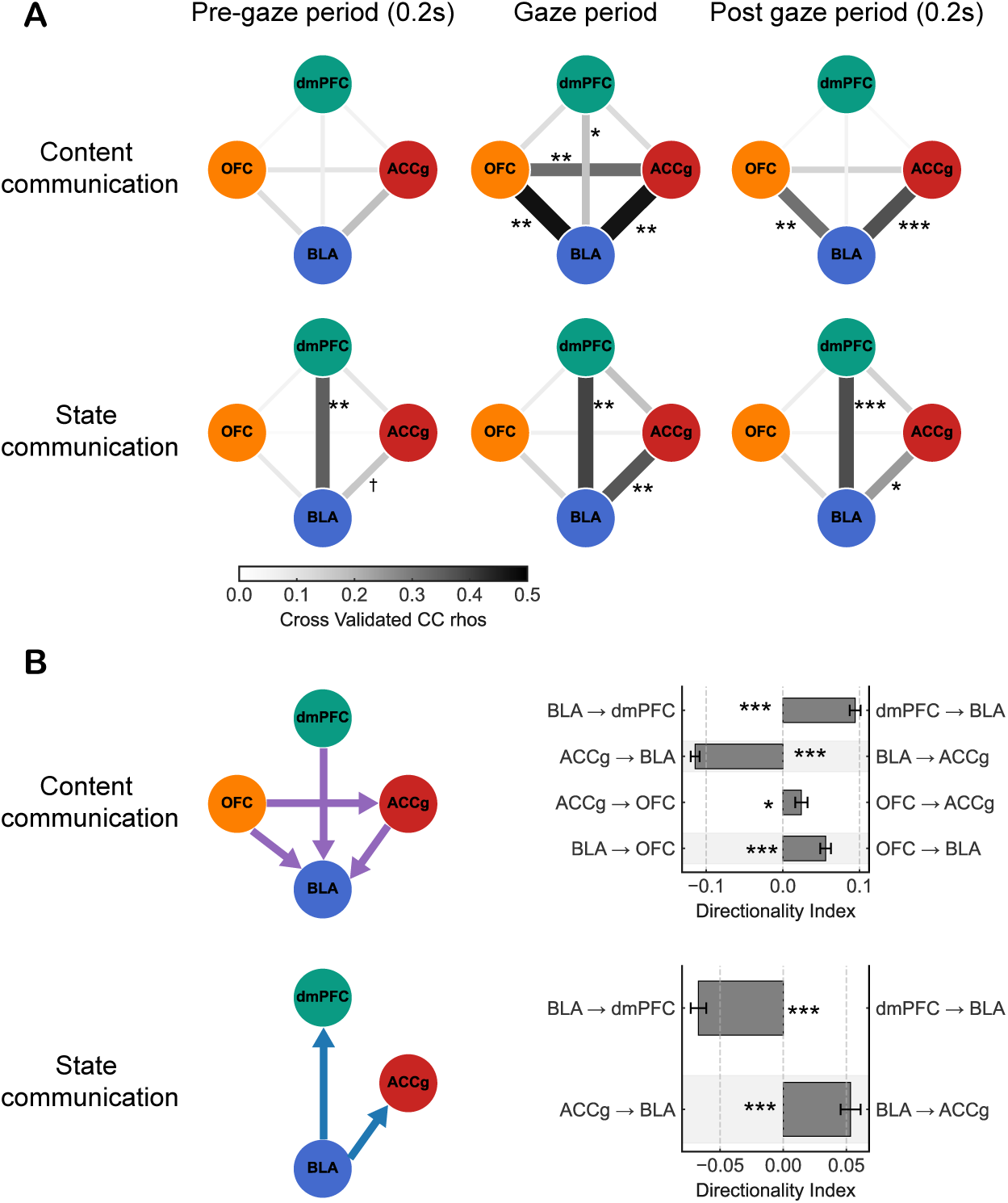
Functional connectivity and directional communication patterns of social gaze primitives in the prefrontal-amygdala circuits. **(A)** Dynamic functional connectivity in the prefrontal-amygdala circuits for gaze content (upper panel) and social state (lower panel) information, respectively, based on zero-lagged CCA. Gaze period: when the eye gaze is on the face or the object. Pre-gaze period: 200 msec before the eye gaze is on the face or the object. Post-gaze period: 200 msec after the eye gaze is on the face or the object. * *P* < 0.05; ** *P* < 0.01; *** *P* < 0.001 with permutation test. **(B)** Directionality of the content (upper panel) and the state (lower panel) information for each pair of brain regions based on CCA lags. Error bar: mean ± SEM. * *P* < 0.05; *** *P* < 0.001 with Wilcoxon signed-rank tests.

We next determined the directionality of functional interactions with respect to content and state information. We performed CCA on each pair of time points using population neural firing rates from pairs of regions (Fig. S14, Fig. S15). If shared neural activity arises at different times across the pairs of regions, we expect to observe a systematic temporal offset. To measure the temporal offset, for each pair of brain regions, we assessed if correlations were stronger for negative or positive lags by calculating a directionality index, which was derived from the sum of correlation coefficients for lags of 200 msec relative to gazing at the target.

For the content information, dmPFC led OFC and BLA; OFC led ACCg and BLA; and ACCg led BLA (Fig. 5B upper panel). By contrast, for the state information, BLA led ACCg and dmPFC (Fig. 5B lower panel). Notably, after removing the linear mixed selectivity cells from these analyses, the functional connectivity between BLA and dmPFC no longer showed significant functional connectivity for the content communication and the directionality for the state communication disappeared (Fig. S16). These results support the critical role of linear mixed selectivity cells in facilitating the communications of social gaze primitives. Taken together, these results suggest that content information flows from dmPFC to BLA, whereas state information flows from BLA to both dmPFC and ACCg.

## Discussion

From behavioral modeling, neural geometry analysis, and communication subspace analysis, we found multiple lines of evidence corroborating that social gaze encoding in the prefrontal-amygdala circuits follows a process akin to compositionality – that is, complex social signals are constructed from a set of simpler, reusable primitives, and crucially, different combinations of these primitives produce distinct meanings ^29–31^. First, behavioral modeling demonstrated that social gaze behaviors emerge from the nonlinear interaction of the three primitives – gaze content, social state, and gaze duration jointly shaped the conspecific’s social gaze behavior. For example, although both high social engagement and face-directed gaze increased social gaze probability, their joint effect was sub-additive, and the influence of content varied depending on fixation duration. This demonstrates that the same primitives, when combined differently, yield different social meanings, consistent with the principle of compositional productivity ^10,13^. This productivity, where slight variations in the combination of primitives (e.g., changing gaze content or social state) result in different behavioral interpretations, is a feature of compositional systems ^13,24^.

Primate social life demands rapid assembly of moment-to-moment gaze cues into coherent signals. Our study decomposed these computations into three primitives and demonstrated that these primitives are not processed in isolation (Fig. 1D). Instead, social gaze interactions critically depended on specific nonlinear combinations of them. Face-directed gaze and high social engagement state each boosted the probability of social gaze, yet their joint impact was sub-additive (Fig. 1E). Further, the effect of face content itself varied with fixation duration (Fig. 1E). Mechanistically, the sub-additive interaction between social engagement state and face-directed gaze suggests a capacity limit or value normalization ^32^ in the receiver: that is, once a monkey is already highly socially engaged, an additional face-directed gaze adds little information, and vice versa. It is important to note that the social state variable was defined based on M1’s own gaze behavior and thus directly reflects M1’s engagement level. The absence of a social-state effect on M2’s reciprocal face-looking behavior therefore does not imply that social state fails to modulate gaze per se, but rather suggests that, in this spontaneous and unconstrained paradigm, one monkey’s engagement state does not necessarily drive the partner’s face-directed gaze on a moment-by-moment basis. We speculate that stronger reciprocity effects may emerge in more structured social tasks that explicitly link gaze behavior to interactive outcomes. Such nonlinear interactions indicate that receivers interpret the combinatorial structure of social gaze primitives rather than relying on any single primitive in isolation. This is consistent with the principles of compositionality, where complex meaning emerges from the combination of simpler, meaningful parts ^10,13^. Compositionality is a hallmark of human language, in which words and syntax combine to express infinite meanings ^30,33^. Similar structure has also been observed in bonobo vocalization ^24^, where call combinations yield novel meanings, and in human facial expressions ^34^, where distinct facial signals combine to modulate the interpretations of speaker’s intent. These parallel findings suggest that, like other communication systems, social gaze may also follow compositional principles, whereby the combination of distinct gaze primitives – such as gaze content, social state, and gaze duration – yields richer and more flexible meanings than any single feature alone.

Second, our study provides evidence that BLA and ACCg abstract social content and social state through a compositional manner (Fig. 2, Fig. 3). Based on the compositionality theory, three kinds of neural relationships between content representation and state representation were tested: (*i*) separability, (*ii*) orthogonality and (*iii*) generalizability. Prediction (*i*) is the minimal prerequisite: if the neural patterns for content and state were identical, a compositional code could not exist. Prediction (*ii*) strengthens this claim: orthogonality is a stricter form of distinctness—orthogonal axes are always distinct, whereas distinct axes are not necessarily orthogonal. Additionally, prediction (*iii*) does not strictly rely on perfect orthogonality, but cross-condition decoding is more robust when the axes approach orthogonality, because it demonstrates the reusable nature of the primitive components across different conditions. Our results suggest that only BLA and ACCg passed all the three predictions. This supports the theoretical prediction that compositional representations require distinct, abstract encodings of primitives that can flexibly generalize across novel combinations ^10–12,14^.

Both content and state signals were widely implemented in BLA, ACCg, dmPFC, and OFC, whereas gaze duration signals were largely absent. Importantly, only BLA and ACCg mapped the content and state onto nearly orthogonal axes and maintained that separation across conditions. Supporting this view, ACCg neurons have been shown to represent social identity even when it is task-irrelevant, consistent with context-independent social coding ^35^. These results are well aligned with the predictions from the compositionality hypothesis ^13,24,29,36^. Linear classifiers trained on one content generalized state to a novel content, and vice-versa, only in these two regions, revealing an abstract, reusable code for both primitives. This orthogonal, generalizable structure is a key signature of factorized representations, where different variables are encoded along independent neural dimensions^37^. The population-level representation composed of multiple gaze primitives should not be interpreted as encoding a single higher-order variable with a fixed meaning. Instead, we propose that it reflects an abstracted summary of the current social significance of a gaze event, integrating information about who is being looked at, the broader engagement context, and temporal structure. Such a representation may support flexible communication across brain networks by allowing downstream regions to access a compact signal that already incorporates multiple dimensions of social gaze, rather than requiring separate integration of individual primitives. Such factorization allows the brain to recombine previous representations in new conditions and therefore supports the generative capacity required for compositional systems. Our findings suggest that BLA and ACCg provide a neural substrate for factorized representation of social gaze primitives, enabling efficient abstraction and reuse of gaze content and social state across varied social scenarios. Furthermore, these findings align with broader theories of cognitive maps, which suggest that the brain encodes diverse information, such as space, value, and social context, using structured, abstract representations that support generalization. Similar principles have been observed in the hippocampus, where neurons form reusable value maps ^38^ and represent individual and group identity across modalities ^39^. Our results suggest that the prefrontal–amygdala circuits may use comparable compositional strategies to guide social gaze.

Third, single-cell analysis uncovered an enrichment of linear mixed selectivity neurons in BLA and ACCg (Fig. 4). This aligns with computational models in which mixed selectivity neurons create high-dimensional, conjunctive representations that preserve the identity of individual primitives while enabling flexible binding, a mechanism essential for compositional encoding ^40–42^. Consistent with theories suggesting that such cells expand representational dimensionality while preserving abstraction ^27^, removing these cells abolished cross-condition generalization in both regions (Fig. 4). Thus, mixed selectivity neurons appear to provide the combinatorial “glue” that binds distinct primitives without letting them interfere. On the other hand, dmPFC and OFC exhibited more limited codes. Both dmPFC and OFC retained the ability to generalize along one dimension – dmPFC for state and OFC for content even though neither region contained a significant proportion of linear mixed-selectivity neurons with respect to the three social gaze primitives. In dmPFC and OFC, the generalization persisted even after removing the few linear mixed selectivity neurons that were present, suggesting that generalization is not solely driven by linear mixed selectivity. Other mechanisms may contribute, such as structured population dynamics that support context-dependent coding ^43^, or projection-specific population codes, where distinct subpopulations encode different variables depending on their downstream targets ^42^. Although dmPFC generalized state but not content, it has previously been linked to abstract representations of agent identity ^44^, suggesting that the coding scheme of dmPFC may be highly context-dependent. Overall, these findings suggest a division of labor in which BLA and ACCg supply a compositional scaffold that other areas tap into selectively. By demonstrating that BLA and ACCg encode social gaze primitives in an orthogonal, generalizable format that leverages linear mixed selectivity, our work reveals a neural grammar that can recombine different social gaze primitives into many social messages. Similar compositional logics may underlie more complex social signals, and their breakdown could help explain social-cognitive dysfunction ^45,46^.

Fourth, content and state information were routed through distinct prefrontal–amygdala channels, providing these associative brain regions a flexible code for social gaze interaction. Communication subspace analysis revealed that content and state are transmitted through separate, time-resolved prefrontal–amygdala pathways: state signals flowed early from the amygdala to the prefrontal cortex, while content signals emerged later and flowed in the reverse direction (Fig. 5). Therefore, the prefrontal–amygdala circuits do not broadcast a single, undifferentiated “gaze signal”. Instead, this anatomically and temporally segregated routing may reflect a neural implementation of compositional structure, where distinct primitives are independently processed and dynamically recombined to support context-dependent social communication ^47^. Additionally, linear mixed selectivity neurons facilitated the communications of content and state information. While our task did not isolate communicative intent, the presence of compositional signatures in spontaneous gaze behavior suggests that this framework may capture core principles of social signal construction, regardless of whether the gaze is explicitly communicative.

State-related population activity was shared early and persistently between BLA ↔ dmPFC and BLA ↔ ACCg, emerging before the animal fixates the target and persisting thereafter (Fig. 5A). By contrast, content-related activity was exchanged later and only during or after the target was viewed. Moreover, content coupling between dmPFC and BLA and between OFC and ACCg faded once the fixation has ended. This finding is consistent with the notion that state information has a longer timescale, whereas content information is deployed on demand for the actual act of looking at a shorter timescale. Furthermore, lagged CCA informed the communication directions of gaze content and social state (Fig. 5B). For state communication, information flowed from BLA to dmPFC and ACCg. However, for content communication, the stream reversed: dmPFC and OFC led both BLA and ACCg, with ACCg leading BLA. Notably, removing linear mixed selectivity neurons eliminated dmPFC–BLA coupling for content and also abolished BLA’s leading role for state communication, implicating these cells as the key components for transmitting primitives from one area to another (Fig. S16). Our results point to a clear division of labor, in which BLA broadcasts a running signal about how socially engaged the animal is, while dmPFC and OFC provide detailed information about what the animal is looking at, perhaps as a feedback-like process. These directional, primitive-specific routings provide a neural network basis for flexibly recombining gaze primitives as social context unfolds.

It is worth noting some limitations of our work. While gaze duration was included as a third primitive, its contribution differed from content and state in both behavior and neural coding. Behaviorally, longer fixations modestly increased the likelihood of the partner’s social gaze, and only with respect to its interaction with content (Fig. 1E): face-directed gazes had extra influence when their gaze durations were short, whereas object-directed gazes mattered more when they were sustained. This crossover effect hints that duration may function as a gain control – prolonged looks may amplify otherwise low-salience cues (objects), whereas short, rapid looks may prevent face cues from saturating the receiver’s response system. None of the four regions showed reliable encoding of duration once content and state were accounted for (Fig. 2). One possibility is that temporal integration occurs in higher-order visual areas that accumulate evidence over hundreds of milliseconds ^48,49^. Social dominance is a fundamental dimension of primate social structure and shapes the interpretation and strategic use of gaze signals. Although dominance relationships were stable across dyads over the course of data collection, our use of fixed partners within each session and a non-chronic recording approach precluded a direct assessment of dominance-related modulation of single-neuron responses. Future studies combining larger social groups, systematic manipulation of hierarchical relationships, or longitudinal recordings across changing dominance contexts will be required to determine how stable social rank interacts with dynamic social gaze signals at the neural level.

Additionally, unlike human language, which has rich context to test the productivity aspect of compositionality ^13,29^, we assessed only two dimensions of interpretation: the probability of partners’ social gaze and their social state in response to different combinations of social gaze. While this provides a first step, it does not fully capture the generative capacity of social gaze signals. Furthermore, because our task did not require explicit communication or cooperation between individuals, the observed gaze patterns may reflect attentional dynamics rather than communicative intent. In this light, not all gaze events are likely to be communicative; many may serve perceptual or exploratory functions. Those that do carry communicative intent may rely on a distinct set of organizing principles, potentially involving a more constrained set of gaze components but greater flexibility in their timing and transitions. Future studies could investigate whether gaze encoding and its compositional structure change in contexts that demand joint action or intentional singling. Such paradigms would be especially informative for testing whether social gaze representations are related to higher-order social cognitive functions such as theory of mind, the ability to infer others’ mental states from gaze and context ^50^. Finally, the gaze primitives identified here may not have fixed meanings on their own. Instead, their meaning may come from how they change and interact over time. This suggests that the ‘grammar’ of social gaze could be dynamic, like tone of voice ^24^ or gesture ^51^, where meaning depends on the flow and timing of signals, not just their individual features.

An important limitation of the present study is that facial expressions were not systematically recorded or quantified. In natural primate social interactions, gaze behavior is often accompanied by facial gestures that can critically shape the perceived emotional valence and communicative meaning of gaze, including whether gaze-based social engagement is interpreted as affiliative, neutral, or threatening ^52^. Because such facial expression signals were not available in the current dataset, our analyses cannot directly address under which circumstances high levels of gaze-based social engagement are interpreted as non-threatening. Accordingly, the social state variable in this study reflects gaze dynamics alone and should not be interpreted as a complete index of social or emotional meaning.

More broadly, gaze behavior is typically embedded within a multimodal social signaling context that includes facial expressions and other bodily cues ^53,54^. In the primate brain, accumulating evidence highlights the importance of contextual information for facial expression and the communicative aspects of bodily gestures ^55,56^. The present study intentionally focuses on decomposing gaze behavior itself into constituent primitives in order to establish how gaze-related variables are represented and combined at the neural population level. As a result, the interdependence between gaze and facial expressions during social interaction is not addressed here. Future studies that simultaneously track gaze, facial expressions, and neural activity will be essential for extending the present framework to a more comprehensive account of naturalistic social communication.

Although the proportion of linear mixed selectivity neurons differed across regions, these differences were modest in absolute magnitude. Accordingly, our interpretation emphasizes relative enrichment rather than large quantitative separation. Future studies with larger samples will be important for refining estimates of regional differences in mixed selectivity prevalence.

Accumulating evidence supports that social information is widely represented in the primate brain ^6,57–60^, with behavioral variables related to social gaze interaction encoded across multiple neural networks ^6,7,61–64^. Future work should explore how compositional processes extend to a broader range of social, behavioral, and emotional responses—including facial expressions, gestures, and homeostatic responses such as body temperature or heart rate, and how primitives from different behavioral domains may combine to generate complex and rich social behaviors.

## Acknowledgments

This work was supported by the National Institute of Mental Health (R01MH110750, R01MH120081, R01MH128190). We are extremely grateful to Ilker Yildirim and Sylvia Blackmore for providing helpful feedback about this work and on the manuscript.

## Author contributions

G.Q. and S.W.C.C. conceptualized the study, O.D.M. and S.F. collected the data, and G.Q. and S.W.C.C. analyzed and wrote the original manuscript. All the authors contributed to reviewing and editing the manuscript.

## Methods

### Experimental Procedures

#### Animals and behavioral task procedures

A total of five rhesus macaques (*Macaca mulatta*) were involved in the study. Neural data were collected from two adult males (M1; Monkey L: 8 years, 15.7 kg; Monkey K: 7 years, 10 kg). The recorded animals (M1) interacted with four partner animals (M2) (three males and one female; Monkeys C, H, L, E; ages 7–8 years), resulting in six distinct social dyads (L–C, L–H, L–E, K–L, K–H, K–E). All monkeys were unrelated, housed in the same colony, and maintained on a 12-hr light/dark cycle with unrestricted food access and controlled fluid access during testing. Procedures adhered to NIH guidelines and were approved by Yale Institutional Animal Care and Use Committee and in compliance with the National Institutes of Health Guide for the Care and Use of Laboratory Animals. No animals were excluded from our analyses.

On each recording session (day), M1 and M2 sat in primate chairs (Precision Engineering, Inc.), 100 cm apart, facing each other (Fig. S1). Eye positions and pupil size were recorded simultaneously from all M1 and M2 monkeys using EyeLink 1000 cameras (SR Research). A standard 5-point screen calibration plus a custom face calibration (using LED markers for eyes, mouth, and face using the face calibration board by the faces of the conspecifics involved; Fig. S1) ^6,19^ were performed before each session, controlled by Psychtoolbox ^65^ and EyeLink toolbox ^66^ in MATLAB (MathWorks). Nonsocial objects (monkey toys) were placed on the M1’s side (Fig. 1B). In a given session, two monkeys spontaneously interacted with gaze in 8-10 consecutive 5-min runs, with a no-vision break between every run (3-min on average) in which they had no visual access to one another. During each social interaction run, middle monitors separating the monkeys were lowered via a remote control, allowing spontaneous face-to-face gaze interactions. During each no-vision break, the middle monitors were raised remotely to block the visual access to one another (Fig. S1). Since some days had only 8 runs, we included the data from the first 8 runs from each day to ensure consistency in data analysis, resulting in a total of 42 sessions (Monkey L: 14 sessions, sessions 1-13, 16; Monkey K: 28 sessions, sessions 14, 15, 17-42) (Fig.1C2).

#### Electrophysiological data collection

Each M1 (recorded monkey) was implanted with a headpost and a recording chamber targeting BLA, dmPFC (Brodmann 8Bm, F6), ACCg (24a, 24b, 32), and OFC (11, 13m) ^67^. MRI and stereotaxic coordinates guided the chamber placements. Single-unit activity was recorded using 16-channel axial array electrodes (U- or V-Probes, Plexon) using a 64-channel Plexon system. Electrodes were lowered via a motorized microdrive (NaN Instruments) through guide tubes, and recordings began after a 30-minute settling period. Analog broadband signals were amplified, band-pass filtered (250 Hz–8 kHz), and digitized (40 kHz) using a Plexon OmniPlex system. All the spiking data were saved for waveform verifications offline and automatically sorted using the MountainSort algorithm ^68^. Only the validated single units were included from each region, without including multi-unit clusters. Neural recordings were simultaneously performed from pairs of regions. In each session, the BLA was recorded together with one of the three prefrontal regions (ACCg, dmPFC, or OFC). Consequently, the BLA was sampled across all sessions, whereas each prefrontal region was sampled in a subset of sessions. In total, we recorded 537 BLA, 236 ACCg, 187 dmPFC, and 241 OFC units across both monkeys (monkey L and monkey K: 225 and 312 BLA cells, 109 and 127 ACCg cells, 92 and 95 dmPFC cells, and 102 and 139 OFC cells, respectively) ^6^. Neural data from each brain region were pooled across recording sessions for analysis, and no analyses required simultaneous recordings across regions.

### Behavioral Data Analysis

#### Social gaze primitives

To study the subcomponents of social gaze, we decomposed social gaze into three primitives for a given gaze bout: gaze content, social state, and gaze duration (Fig. 1A). Each gaze event was defined as when M1 looked at M2’s face (face bout) or object (object bout). From each day’s calibration, the face region of interest (ROI) was defined by the four corners of a monkey’s face, with the object ROI having the same surface area as the face ROI. Fixations were identified using EyeMMV toolbox ^69^ implemented in MATLAB. We detected fixations using spatial and duration parameters. Specifically, we used t1 = 1.18 and t2 = 0.59 degrees of visual angle for the spatial tolerances, and a minimum continuous duration of 70 msec.

In this study, we distinguish between event-level gaze features and contextual variables that characterize the interactional background in which individual gaze events occur. Gaze content and gaze duration are defined at the level of individual gaze bouts, whereas social state is a contextual variable that reflects a sustained mode of social engagement across multiple gaze events. Accordingly, frequency-based measures are used here as an operational estimate of this contextual social state, rather than as a primitive defined at the level of individual gaze events.

For each gaze bout, social states were defined as each monkey exhibiting either high social engagement (high) or low social engagement (low). This analysis was done for each M1 separately (Fig.1C2). The high social engagement was operationally defined as M1 being more likely to look at M2’s face relative to the object, while the low social engagement was defined as M1 being more likely to look at the object relative to M2’s face. To calculate the social states at a resolution for capturing natural fluctuations, we first divided each of the eight 5-min runs into three segments (100-sec each) (Fig. 1C1). We selected 100-sec segments as it provided the highest temporal resolution while still reliably capturing state-like dynamics. For each segment, we calculated the social state index as the normalized fraction of face bouts as (Eq. 1):

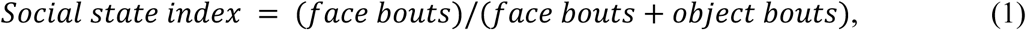

where face bouts and object bouts were defined as the number of face and object bouts during each segment, respectively (Eq. 1). To categorize social state as a binary variable for further analysis, each segment was labeled as either having a high or a low state depending on whether the social state index was above or below the median value of these indices within that session. Similarly, for the gaze duration, each gaze bout was labeled as either having a long or a short duration depending on whether its duration was above or below the median value of the gaze durations within that session. Finally, the gaze content reflected whether the monkeys were looking at the partner’s face or the nonsocial object. Although social state is derived from gaze content at the behavioral measurement level, the aim of this study is to examine whether contextual variables (state) and event-level variables (content) are represented in dissociable formats at the neural population level.

#### Conceptual grounding of the three gaze primitives

The choice of gaze content, gaze duration, and social state as candidate gaze primitives is grounded in prior behavioral, neuroethological, and social neuroscience literature:

i. Gaze content (“who or what is being looked at”) is a foundational variable in social attention research across species, distinguishing social from non-social targets and modulating neural responses in amygdala and prefrontal circuits ^2,6,7,18^.
ii. Gaze duration captures the temporal structure of individual gaze events. Duration is known to modulate communicative meaning in gaze behavior, including signaling intent, threat, or affiliation, and has long been treated as a separable dimension of gaze behavior in both primate and human studies ^2,70,71^. In developmental and comparative psychology traditions, gaze duration has been long studied to examine violation of expectation ^72^.
iii. Social state captures the contextual background of an interaction, specifically the sustained level of social engagement across multiple gaze events. Contextual or state-like variables are widely used in social neuroscience to describe slow-varying internal or interactional modes—such as social engagement, vigilance, or arousal—that modulate the interpretation of momentary actions ^73–75^. Critically, such variables are not attributes of single events, but characterize the broader interactional context in which events occur.

Importantly, we do not claim these primitives are exhaustive. Rather, they constitute a minimal set of event-level (content, duration) and context-level (state) dimensions at a common level of abstraction that enables (i) tractable behavioral tests of whether different components contribute distinct predictive power, and (ii) corresponding neural tests of whether these components are represented dissociably and in an abstract format.

#### Pupil size

To examine the relationship between pupil size and social engagement state, we computed the Pearson correlation between pupil size and the social state index. Pupil size was recorded from the monkeys using EyeLink 1000 cameras (EyeLink 1000, SR Research). Pupil size data were min-max normalized with a [0, 1] range using a 100-Hz sampling rate. To align with the segment structure used for defining state variables, we examined the mean pupil size from each segment.

#### General linear mixed-effects modeling of M2’s social gaze

To test how different M1’s social gaze primitives and their combinations influence M2’s social gaze behavior, we modeled the likelihood of M2’s social gaze using a generalized linear mixed-effects model (GLMM) with a binomial distribution and logit link function, implemented using the ‘lme4’ package in R (Eq. 2). Fixed effects included M1 content (Content), M1 state (State), and M1 duration (Duration), as well as all their two-way and three-way interactions. Random intercepts were included for session, M1 identity, and M2 identity, to account for repeated measurements and the hierarchical structure of the data. The model was defined as (Eq. 2):

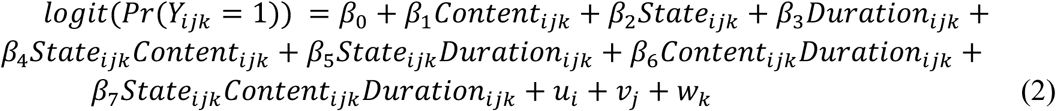

where *Y_ijk_* denotes the binary outcome of M2’s social gaze (1 means M2 gazed at M1’s face within 1 sec after M1’s gaze event ended; 0 means M2 did not gaze at M1’s face within 1 sec after M1’s gaze event ended) for an observation made on session *i*, with M1 participant *j* and M2 participant *k*; and *u_i_*, *v_j_*, *w_k_* ∼ N(0, σ²) represent random intercepts for sessions, M1, and M2, respectively.

Fixed effects were evaluated using Type III Wald chi-square tests via the ‘car’ package in R. Post hoc pairwise comparisons were conducted on the estimated marginal means (EMMs) using the ‘emmeans’ package in R. Wald z-tests were applied to test the significance of the estimated differences between conditions. Importantly, the standard errors of these differences account for the covariance structure of the mixed-effects model, including random effects. Multiple comparisons were corrected using the Tukey HSD adjustment. Notably, the reported 95% confidence intervals for marginal means represent uncertainty around the group-level predictions, whereas the significance tests rely on the precision of the estimated differences; thus, statistical significance may be detected even when confidence intervals overlap visually.

In a complementary analysis, we performed an additional GLMM to assess how M1’s social gaze primitives influenced M2’s social state. The M2’s social state was defined using the same methods as previously described for M1. Because M2 did not have a physical object, however, we used the matching area corresponding to M1’s object zone from M2’s perspective as the nonsocial reference region. The same fixed and random effect structure was used, with the binary outcome defined as whether M2 was in a high versus low social engagement state within a 100-sec window following M1’s gaze event.

##### Control analyses for dominance hierarchy

### Dominance as an additional predictor in the GLMM

To assess whether stable dominance–subordination relationships between monkeys could account for the observed social gaze coordination effects, we extended the original generalized linear mixed-effects model (GLMM; Eq. 2) to explicitly include partner dominance as an additional predictor of M2’s social gaze behavior.

Each M1–M2 dyad was assigned a binary dominance label based on consistent behavioral observations across recording sessions (dominant = 1 if M1 was dominant over M2; 0 otherwise). Dominance assignments were independently evaluated by two observers familiar with the animals and colony during the period of data collection. Importantly, this dominance variable reflects a stable, pair-level social hierarchy and does not vary on a trial-by-trial basis.

Using the same dataset and preprocessing pipeline as in the primary behavioral analyses, we fit an extended GLMM with a binomial distribution and logit link function to predict the probability of M2’s social gaze following each M1 gaze event (Eq. 3):

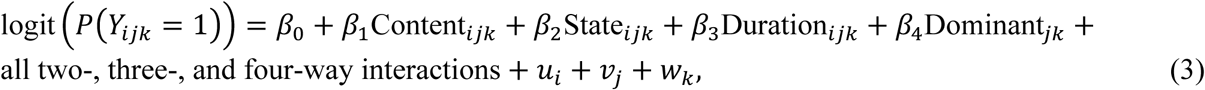

where *Y_ijk_* denotes whether M2 gazed at M1’s face within 1 s after the end of M1’s gaze event in session i, with M1 identity j and M2 identity k. Content (Face vs Object), State (High vs Low), and Duration (Long vs Short) were defined identically to the original GLMM analysis. Random intercepts *u_i_*, *v_j_*, and *w_k_* were included for session/day, M1 identity, and M2 identity, respectively, to account for repeated measurements and the hierarchical structure of the data. Models were implemented in R using the lme4 package (glmer, Laplace approximation).

Fixed effects were evaluated using Type III Wald chi-square tests implemented via the car package, consistent with the original analysis pipeline. Significance was assessed at α = 0.05. Estimated fixed-effect coefficients are reported as log-odds with 95% confidence intervals.

### Alternative specification: dominance as a random effect

As an additional robustness check, we repeated the analysis using an alternative model in which dominance was treated as a random intercept rather than a fixed effect. In this specification, random intercepts were included for session/day, M1 identity, M2 identity, and dominance, while the fixed-effect structure retained Content, State, Duration, and their interactions. This complementary model tested whether allowing dominance-related variance to be absorbed at the group level altered the estimated effects of the gaze primitives.

### Neural Data Analysis

#### Preprocessing of single-unit data

To estimate continuous time-dependent firing rates, timestamps of spiking events were resampled at 1 KHz and converted into binary spikes. Spike trains were then convolved with a symmetric Hann kernel (sample rate: 100 Hz; kernel width: 0.3 sec) in MATLAB. The resulting firing rates were then min–max normalized to the [0, 1] range. The peri-stimulus time histograms (PSTHs) were then calculated for three periods of interest around each gaze bout: (i) the pre-gaze period (200 msec before the gaze event to the face or object), (ii) the gaze period (during the gaze event), (iii) the post-gaze period (200 msec after the end of the gaze event). We used the averaged firing rate during these three periods for further analysis. All neural analyses were time-locked to individual gaze events. A gaze event was defined as a continuous fixation to a specific region of interest (ROI; face or object), as determined by fixation detection and ROI assignment procedures described above. Gaze events varied naturally in duration. For each gaze event, neural activity was quantified as the average firing rate over the full duration of that event. Importantly, neural activity was not rescaled or truncated to a fixed duration across events; instead, each event contributed a single observation reflecting its own temporal extent.

#### Encoding model of representational dissimilarity matrix (RDM)

To investigate how social gaze primitives are encoded by the four neural populations, we applied a general linear model (GLM) to characterize the representational structure of neural population activity during each gaze event (i.e., gaze period). We examined whether three primitives – gaze content, social state, and gaze duration – could explain neural dissimilarity patterns across conditions (see below). Each of these primitives was represented as a representational dissimilarity matrix (RDM), resulting in three hypothetical RDMs that were predictors in the regression analysis (Fig. 2A).

To construct the RDMs of each brain region, we began by categorizing each gaze event according to the three primitives, yielding eight unique event conditions ([content/state/duration]: face/high/long, object/high/long, face/high/short, object/high/short, face/low/long, object/low/long, face/low/short, and object/low/short). For each neuron, we computed its average firing rate in the gaze period on a gaze event-by-gaze event basis. These averaged firing rates were saved in a 3D array, with dimensions corresponding to neuron, condition, and gaze event number. Neural activity was analyzed separately for each brain region of interest (ROI), including BLA, ACCg, dmPFC, and OFC. To generate a neural RDM for each brain region, we first averaged each neuron’s firing rate across all gaze events within each condition, resulting in one condition-level population vector per ROI. Pairwise Euclidean distances were then computed between these eight vectors to form an 8 × 8 neural RDM. These dissimilarity matrices were normalized to a [0, 1] range. These neural RDMs were used to visualize the representational geometry of each brain area by projecting them onto the first three dimensions using multidimensional scaling (MDS) (Fig. 3D). To quantify how well the low-dimensional MDS embeddings preserved the original high-dimensional neural distance structure, we computed the explained variance as the squared Pearson correlation (r²) between pairwise distances in the original neural RDM and those in the corresponding 3D MDS embedding.

To evaluate the contribution of each primitive, we applied multiple linear regression. The upper triangular values (excluding the diagonal) of the neural RDM were vectorized and used as the dependent variable. The three hypothetical model RDMs (for content, state, and duration) (Fig. 2A) were also vectorized as independent variables. The regression model was specified as (Eq. 4):

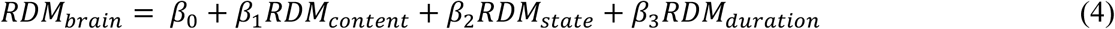

We used ordinary least squares regression to estimate the beta coefficients, allowing us to quantify how strongly each variable contributed to the neural dissimilarity structure during the gaze time window.

To assess statistical significance, we conducted a permutation-based shuffle analysis. For each brain region, the condition labels in the neural data were randomly shuffled 1,000 times while keeping the structure of the hypothetical RDMs fixed. For each shuffled dataset, we recomputed the neural RDM and re-ran the GLM, generating null distributions of the regression coefficients. Statistical significance for permutation-based analyses was then assessed using a two-tailed permutation test defined in terms of the absolute value of the test statistic. For an observed statistic *β_obs_* (e.g., a regression coefficient) and a null distribution 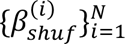 obtained from *N* label-shuffled permutations, the permutation p value was computed as (Eq. 5):

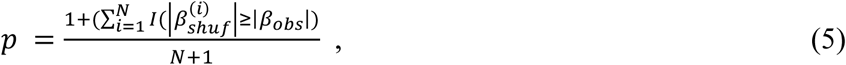

where I(⋅) denotes the indicator function. This formulation evaluates extremeness in either direction relative to the permutation null distribution and corresponds to a two-tailed test.

To further assess the unique contribution of each primitive, we performed a partial correlation analysis in which the neural RDM was correlated with each model RDM while controlling for the remaining primitives (Fig. S7). Partial correlations were computed by regressing out the other model RDMs from both variables and correlating the resulting residuals. This analysis provides a scale-invariant complement to the GLM results and yields equivalent inferences regarding the unique contribution of each primitive.

#### Population decoding analyses

##### Content decoding

For population decoding of content, we used linear Support Vector Machine (SVM) classifiers implemented from the scikit-learn toolbox ^76^. All the recorded neurons were included, regardless of their content selectivity. The classifiers were trained to distinguish between face and object content using neural population activity recorded during individual gaze events from each brain area, treating each event as an observation for decoding. To construct a pseudo-population, data from all recording sessions were pooled, and only neurons with more than 10 gaze events for each content condition were included. For each neuron, exactly 10 gaze events were randomly sampled from the face condition and 10 gaze events from the object condition, ensuring perfectly balanced class sizes at the event level. A 5-fold cross-validation was used, where the dataset was randomly divided into five subsets, with the classifier trained on four subsets and tested on the remaining one subset. This was repeated five times to calculate average decoding accuracy. The entire cross-validation procedure was repeated 1,000 times, and the final decoding accuracy was computed as the mean across all repetitions. Statistical significance was assessed by comparing this mean accuracy to a null distribution generated from label-shuffled data. Decoding was considered significant if the mean accuracy exceeded the 95th percentile of the null distribution. All subsequent decoding and generalization analyses in this study used the same procedure, with exactly 10 gaze events per condition per neuron and identical cross-validation and shuffling controls, unless otherwise specified.

##### State decoding

Neurons with more than 10 gaze events in each state group (high and low state groups, same as for the content decoding analysis described above) were selected for the state decoding. The decoding procedure was the same as described for the content decoding procedure, except that the gaze events were sorted and labeled by high and low state groups. Statistical significance was assessed by comparing this mean accuracy to a null distribution generated from label-shuffled data. Decoding was considered significant if the mean accuracy exceeded the 95th percentile of the null distribution.

##### Cross-primitives decoding

To assess whether the neural representation of content is shared with state, we performed cross-primitives decoding (Fig. S8B). If the representations of these variables are not shared (see below for the additional orthogonality test), we expect the cross-primitives decoding to be poor. Specifically, we trained a linear SVM classifier to decode content (face vs. object) using only gaze events labeled by content and then tested the classifier’s ability to decode state (high vs. low) using the state-labeled gaze events, and vice versa. All procedures were the same as those described for the content and state decoding analyses. A 5-fold cross-validation approach was used, repeated 1,000 times. Only neurons with at least 10 gaze events in each relevant condition were included. Statistical significance was determined by comparing the observed decoding accuracy to a null distribution generated from label-shuffled data, with decoding considered significant if accuracy exceeded the 95th percentile of the null distribution.

##### Cross-conditions decoding

To examine whether the neural representation of state or content is structured in an abstract format and can be generalized to new conditions (e.g., a linear decoder for state in face-directed events can be used in unseen object-directed events), we performed a cross-condition generalization analysis of social gaze primitives (distinct from the cross-primitives decoding described before) (Fig. S8D). The procedure followed the same approach as described for the content and state decoding analyses, using linear SVM classifiers with 5-fold cross-validation repeated 1,000 times. For each neuron, we trained the classifier to decode one variable (e.g., content: face vs. object) using neural activity from single gaze events that occurred under one condition (e.g., high state) and tested the classifier on gaze events for the other condition (e.g., low state). The average decoding accuracy across both state conditions (i.e., training on high state and testing on low state, and vice versa) was defined as the content generalization performance. The same procedure was used to compute state generalization performance across content conditions (i.e., training on face and testing on object, and vice versa). Only neurons with more than 10 gaze events in each relevant condition were included. Statistical significance was assessed by comparing decoding accuracy to a null distribution generated from label-shuffled data, with significance determined by exceeding the 95th percentile of the null distribution.

#### Neural state space analysis

To test the orthogonality between content and state information representations, we computed condition-specific population vectors and quantified the geometry of their representational structure using neural state space analyses (Fig. S8C) ^26^. Neural spiking data were organized as a 3D matrix of shape (neurons × conditions × gaze events), with four sub-conditions defined by a 2×2 factorial design: content (face vs. object) and state (high vs. low). For each neuron and condition, the mean firing rate was computed for different gaze events, yielding four condition-specific population activity vectors per region (content/state): face/high, object/high, face/low, and object/low. Two encoding axes were defined to represent neural selectivity for content and social state. The content axis was defined as the mean firing rate difference between face and object conditions, averaged across state levels (Eq. 6):

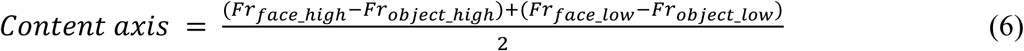

The state axis was defined as the mean firing rate difference between high and low state conditions, averaged across content levels (Eq. 7):

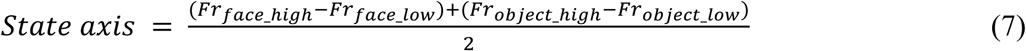

The angle between these axes was calculated using subspace angles from the SciPy library, which computes the principal angle between subspaces. The resulting angle (in degrees) quantifies the degree of independence between content and state representations within the neural population. To assess the statistical significance of the observed angle, a permutation test (1,000 iterations) was used. On each iteration, condition labels were randomly permuted across conditions within each neuron. The content and state axes were then re-estimated, and the angle between them was calculated. The permutation p-value was defined as the proportion of iterations in which the permuted angle was smaller than or equal to the observed angle.

To visualize the structure of neural representations in a reduced dimensional space, principal component analysis (PCA) was applied to the matrix of condition vectors after mean-centering across conditions. The first two principal components were used to project the condition vectors and the content/state axes. Projected content and state vectors were normalized to unit length and plotted as arrows originating from the origin.

#### Shattering dimensionality analysis

To examine the neural separability among four kinds of gaze events or sub-conditions (face gaze in high state, object gaze in high state, face gaze in low state, and object gaze in low state), we conducted a decoding-based shattering dimensionality (SD) analysis ^25^. For each combination of two kinds of gaze events (in total six combinations from four sub-conditions), the decoding procedure followed the same approach as described for the content and state decoding analyses, using linear SVM classifiers with 5-fold cross-validation repeated 1,000 times. The averaged accuracy from six decoding procedures was reported as SD. The larger value means that sub-conditions in that brain region are more separable. Statistical significance was assessed by comparing decoding accuracy to a null distribution generated from label-shuffled data by the same decoding procedure, with significance determined by exceeding the 95th percentile of the null distribution. Shattering dimensionality was estimated by averaging decoding performance across all possible pairwise condition partitions, providing a population-level measure of representational flexibility rather than pair-specific decoding strength.

#### Single-cell classification

To determine the content or state selectivity of individual cells (Fig. 4A), we first performed a linear regression on averaged firing rates during gaze time using the model (FR ∼ content + state) for each cell, testing for main effects of content and state (Fig. 4A). Duration was not included because the encoding model (Eq. 4) confirmed that it cannot be significantly represented in all four brain areas. Based on the resulting p-values, neurons were classified into different categories.

If *P*(content) < 0.05 and *P*(state) ≥ 0.05, the neuron was classified as a pure content-selective cell. If *P*(state) < 0.05 and *P*(content) ≥ 0.05, the neuron was classified as a pure state-selective cell. If both *P*(content) < 0.05 and *P*(state) < 0.05, the neuron was classified as a candidate mixed selectivity cell. To further distinguish between linear and non-linear forms of mixed selectivity, we compared the initial linear regression model to an extended regression model that included the interaction term using a nested model comparison approach. Cells with significant interaction effects (*P*(interaction) < 0.05) were designated as having non-linear mixed selectivity. Cells without a significant interaction effect (*P*(interaction) ≥ 0.05) were classified as showing linear mixed selectivity. This procedure resulted in four categories:

i. Pure content selectivity: significant main effect of content only.
ii. Pure state selectivity: significant main effect of state only.
iii. Linear mixed selectivity: significant main effects of both state and content, but no significant interaction.
iv. Non-linear mixed selectivity: significant interaction between state and content.

#### Population PSTHs recoded by state or content preference

To characterize the temporal dynamics of population selectivity for gaze content and social state, we computed population peristimulus time histograms (PSTHs) aligned to gaze onset. Analyses were performed separately for each brain area and for each gaze primitive (content or state). Neurons were included if they showed significant modulation by the corresponding primitive in the single-neuron classification analysis (pure selective or mixed selectivity neurons).

For each neuron, trials from the two conditions (face vs object for content; high vs low social state for state) were randomly split into independent training and test sets with balanced trial counts. Neuronal preference was defined on the training set as the condition eliciting the higher mean firing rate in a post-gaze window (0–200 ms relative to gaze onset). Population PSTHs were then computed from held-out test trials only to avoid circularity. Firing rates were aligned to gaze onset, min–max normalized to the [0, 1] range within each neuron, and averaged across neurons within each area.

Statistical significance of preferred versus non-preferred differences over time was assessed using a cluster-based permutation test on the neuron-wise difference time courses (preferred minus non-preferred). A one-sample t-test against zero was computed at each time bin, clusters were defined as temporally contiguous bins exceeding a fixed t-threshold (*P* < 0.001), and cluster significance was determined relative to a permutation-based null distribution generated by sign-flipping across neurons. Only clusters spanning at least 100 ms were considered significant. Significant time windows are indicated by black horizontal bars in the PSTH plots.

#### Control analysis for visual stimulation history

To test whether neural modulation attributed to social state could be explained by short-term visual stimulation history, we performed a single-neuron control analysis explicitly modeling the effect of the immediately preceding gaze event.

For this analysis, history was defined as the category of the immediately preceding gaze event (face = 1, object = 0), state was defined as the current social state (high = 1, low = 0), and firing rate (FR) was defined as the mean neural response during the current gaze event (post-gaze window), consistent with all other single-neuron analyses in the manuscript.

Using the same event-aligned firing-rate measure employed elsewhere, we constructed a long-format dataset containing, for each neuron and gaze event, the firing rate, state, and history. For each neuron, we fit an ordinary least squares regression model of the form (Eq. 8):

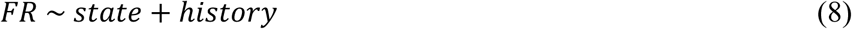

Neurons were retained only if all four state × history combinations were observed, ensuring model identifiability. For each neuron, we extracted the regression coefficients (β_state, β_history) and their associated p-values. Analyses were performed separately for each brain region.

To summarize the population-level structure of effects, we performed three complementary analyses within each region:

i. we quantified the fraction of neurons significantly modulated by state, by history, and by both predictors (*P* < 0.05);
ii. we tested whether the overlap between state-significant and history-significant neurons exceeded chance using a hypergeometric test (probability of observing ≥ the measured overlap given marginal counts); and
iii. within neurons significant for both predictors, we assessed the relationship between β_state and β_history using Pearson correlation.

#### Canonical Correlation Analysis (CCA)

To test how content and state information flowed through the prefrontal-amygdala circuit, we applied canonical correlation analysis (CCA) ^77,78^ to examine the relationships between population activity in BLA, ACCg, dmPFC, and OFC. First, spiking activity from any two brain regions was aligned based on matching behavioral conditions – specifically, face vs. object for content information communication, and high vs. low state for state information communication. For content information analysis, trials were aligned such that face events in one region corresponded to face events in the other region, and similarly for object events; however, state information was not aligned. Conversely, for state information analysis, high and low state trials were aligned across regions, while content (face vs. object) was not aligned. We then performed a PCA across gaze events to reduce the dimensionality and to obtain the first 10 PCs for each brain region. The gaze events were then divided into 5 equal parts (training set and testing set) for cross-validation (5-fold cross-validation). The PCs of the training set of each brain region were used to perform canonical correlation analysis (CCA) to obtain the first pair of canonical correlation components (L2 regularization, λ=0.5). The PCs of the testing set from each brain region were then projected onto the first pair of canonical correlation components, and the correlation was determined by the Pearson correlation coefficient between these projections from each region. This analysis was performed for each pair of three periods (pre-gaze, gaze, and post-gaze) to generate a 3×3 cross-validated correlation coefficient matrix (Figs. S14-S15). For each iteration, 50 gaze events per condition (i.e., face and object gaze events for content communication; high and low state gaze events for state communication) were randomly selected from each brain region using bootstrapping. This process was repeated 1,000 times, and the resulting correlation matrices were averaged to produce the final heatmap. To test the functional connectivity between each pair of brain areas, gaze event labels within each region were independently shuffled to disrupt the condition alignment between regions. The correlation analysis described above was then repeated 1,000 times to generate a null distribution. Functional connectivity between two regions was considered significant if the mean Pearson correlation coefficient exceeded the 95th percentile of the null distribution.

To quantify the lead-lag relationship in information exchange between brain regions, we computed a directionality index defined as (Eq. 9):

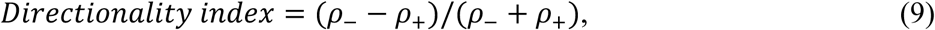

where ρ₋ and ρ₊ denote the sum of min-max normalized correlation coefficients at negative and positive time lags, respectively ^79^. A positive index (from region A to region B) indicates that region A leads the region B in directional information flow, whereas a negative index suggests that region B leads region A. To assess statistical significance, we tested whether the directionality index of each pair of brain areas is significantly different from 0 with a Wilcoxon signed-rank test.

#### General notes on statistics

Overall, all statistical analyses were performed using MATLAB (MathWorks), Python (scikit-learn, statsmodels), and R (lme4, car, emmeans). Two-tailed tests were used unless otherwise noted. Significance thresholds were set at *P* < 0.05. Permutation testing was used to assess significance for model fits and decoding performance. For each test, 1,000 permutations were performed to generate null distributions, and empirical p-values were calculated accordingly. Cross-validation procedures and data shuffling methods are described in detail above in the respective sections.

## Data Availability

Behavioral and neural data presented in this paper and the analysis codes will be available through https://github.com/changlabneuro/social_gaze_compositionality upon acceptance of the manuscript. Source Data are provided with this paper.

## Code Availability

Behavioral and neural analysis codes will be available through https://github.com/changlabneuro/social_gaze_compositionality upon acceptance of the manuscript.

## Supplemental Figures and Figure Legends

**Figure S1.**
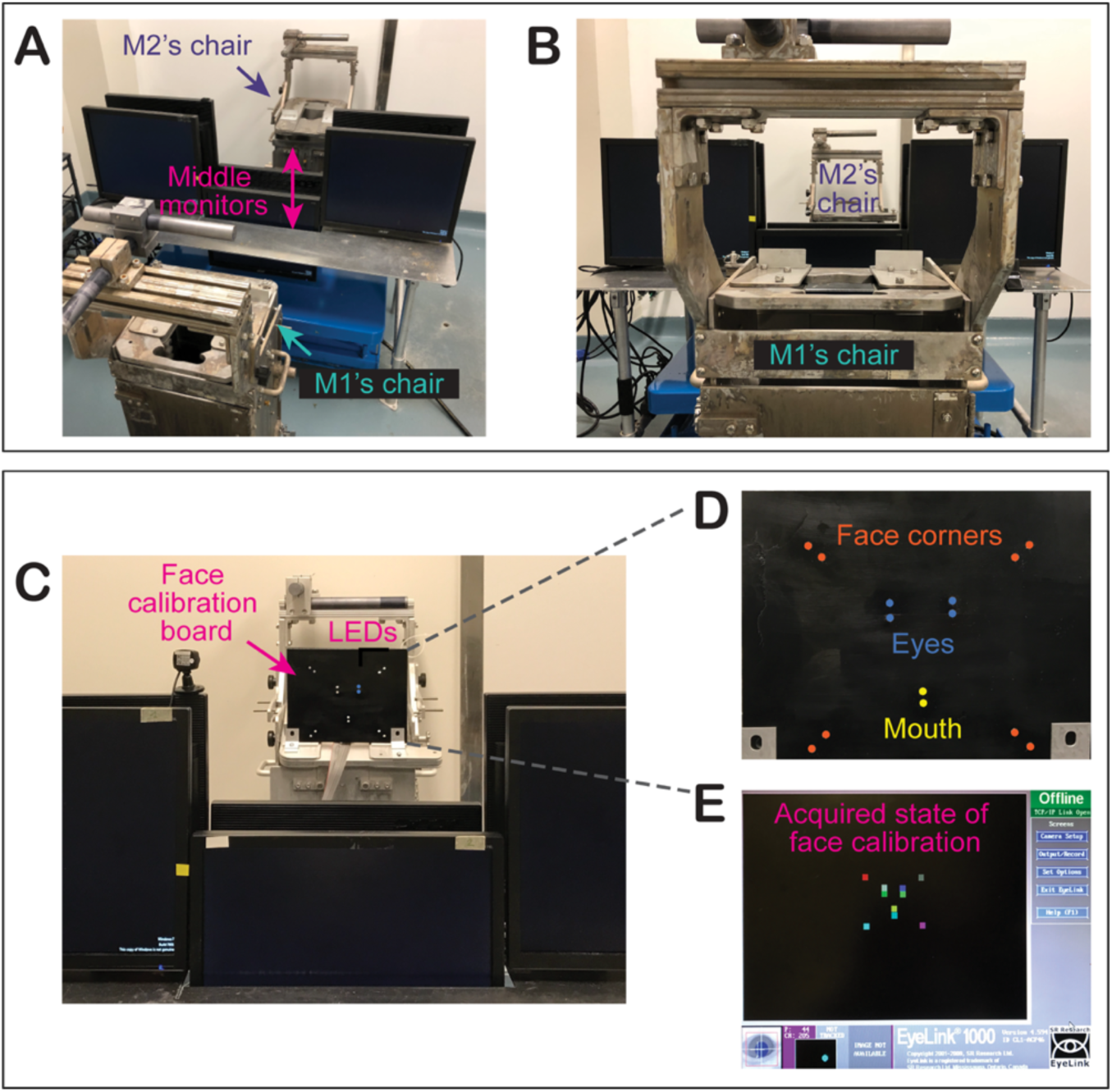
Experimental setup and face calibration. Related to Fig. 1B. **(A–B)** Setup for real-life social gaze interaction, with two monkeys positioned face-to-face. The central monitors could be remotely raised or lowered via a hydraulic system to control visual access. **(C–D)** Custom LED face calibration board with lights aligned to key facial landmarks (eyes, mouth, corners). Two versions fit different face sizes. LED colors in **(D)** are illustrative; actual lights used a single color. **(E)** An example calibration result showing gaze-to-LED position mapping (colored squares).

**Figure S2.**
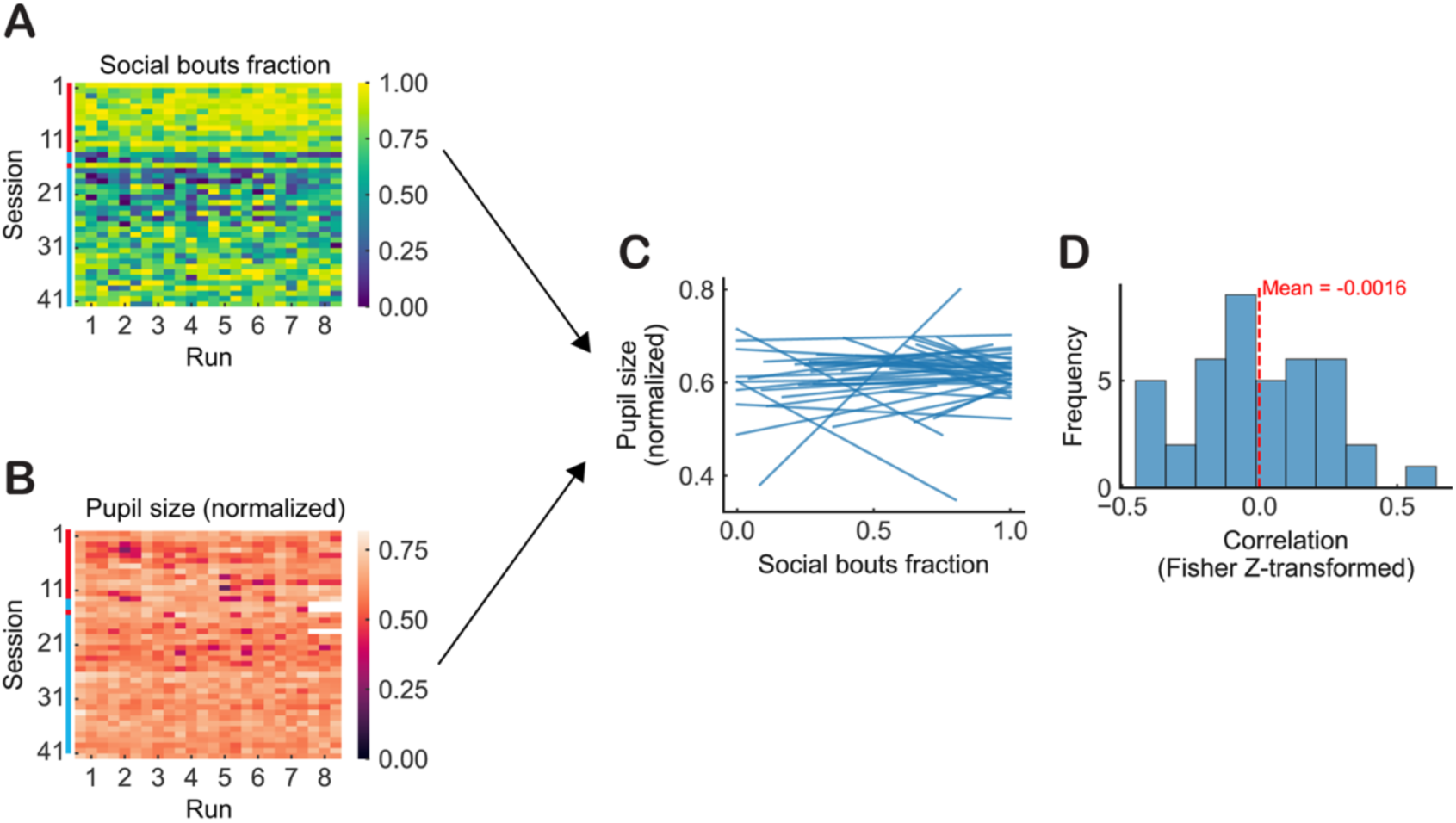
Correlation between pupil size and social state. Related to Fig. 1C. **(A)** Normalized social gaze (face) bouts fraction across runs and sessions (red marks: sessions 1–13 and 16 from Monkey L; cyan marks: sessions 14–15 and 17–42 from Monkey K). **(B)** Normalized pupil size (min-max normalized to the [0, 1] range within each session) across runs and sessions (red marks: sessions 1–13 and 16 from Monkey L; cyan marks: sessions 14–15 and 17–42 from Monkey K). Note that the white segment in the pupil size matrix means the pupil data was missing. **(C)** Line plots showing the relationship between normalized pupil size and the fraction of social gaze bouts across runs within each session. Each line represents a single session. **(D)** Histogram of session-wise Pearson correlation coefficients (Fisher Z-transformed) between pupil size and social bout fraction. The red dashed line marks the mean correlation (−0.0016), indicating no systematic relationship between pupil size and social engagement across sessions (*P* = 0.97, one sample t-test with 0).

**Figure S3.**
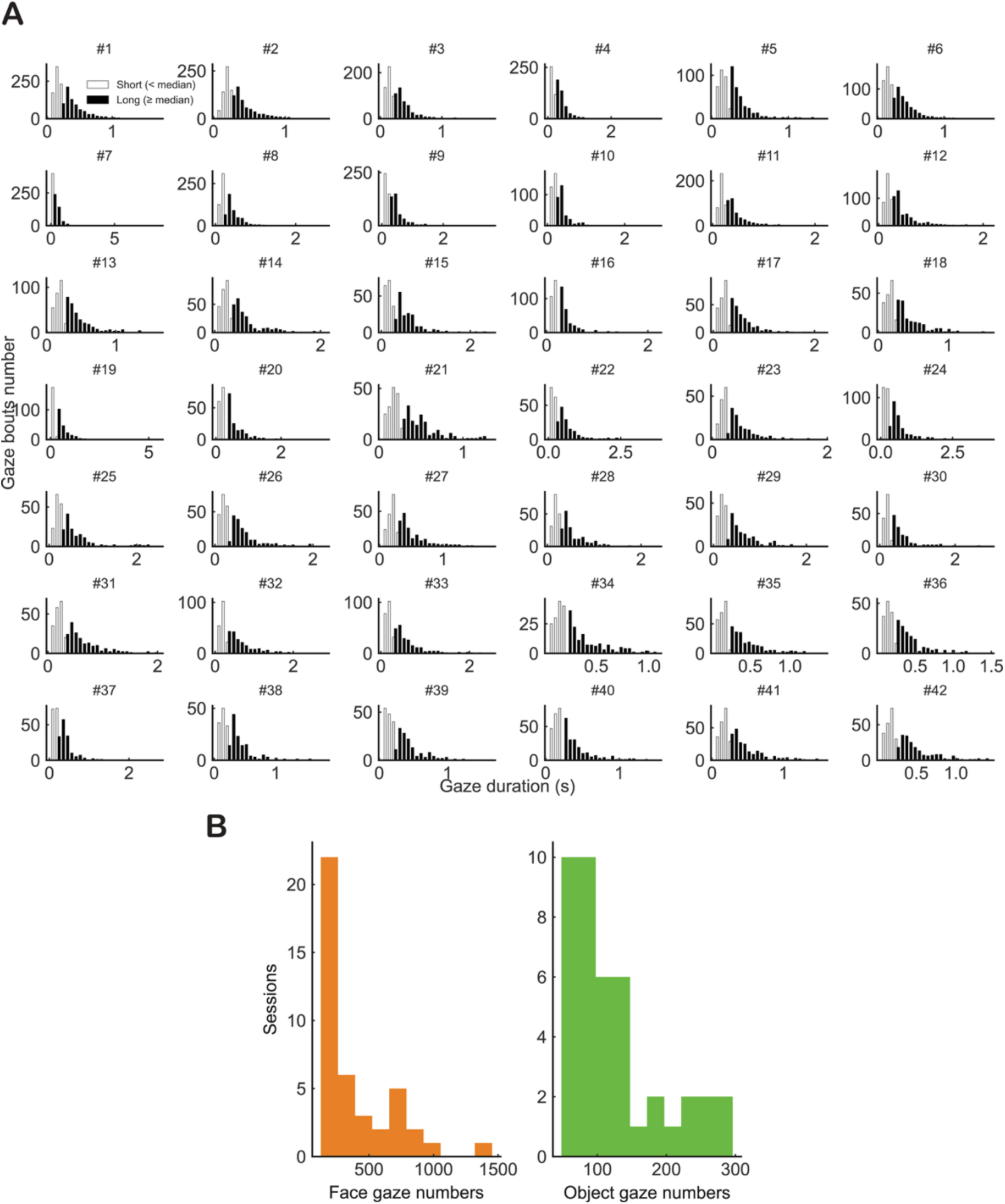
Distributions of gaze duration and gaze content. Related to Fig. 1. **(A)** The distributions of gaze durations from all 42 sessions by session. Each gaze bout was labeled as a short (open bar) or long (filled bar) duration event based on whether its duration was lower or higher than the median of duration in a given session. **(B)** Distributions of gaze content across all 42 sessions.

**Figure S4.**
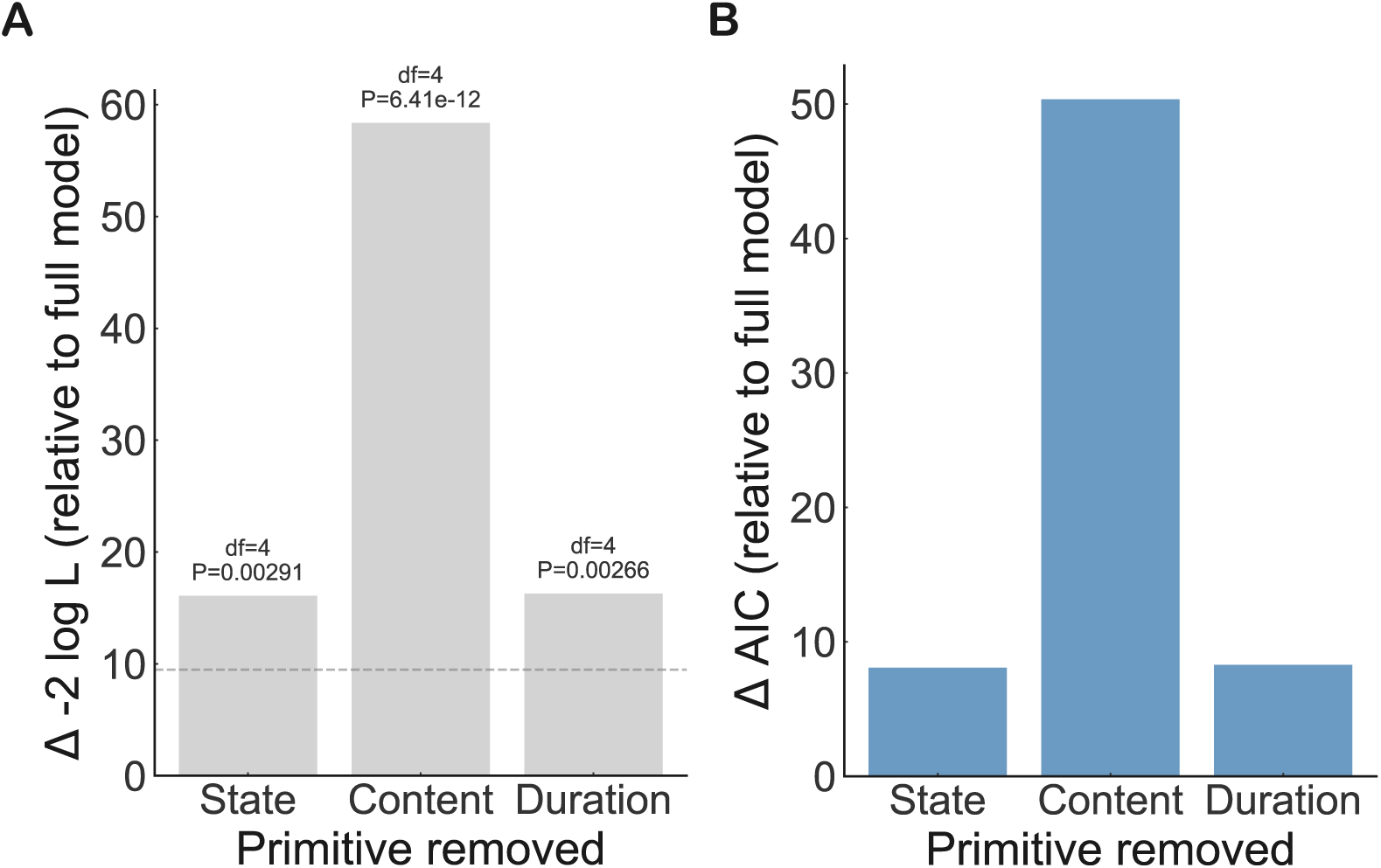
Behavioral validation of gaze primitives and compositional structure. **(A)** Block-wise likelihood-ratio tests (Δ−2 log L) relative to the full model demonstrate that removing gaze content, social state, or gaze duration each significantly degrades model fit, indicating that all three primitives contribute non-redundant explanatory information. P values are from χ² tests (df = 4). The dashed line indicates the χ² critical value for α = 0.05, used to assess statistical significance in these block-wise likelihood-ratio tests. **(B)** ΔAIC relative to the full model when each gaze primitive is removed (along with its associated interaction terms). Removing any primitive substantially worsens model performance, with gaze content producing the largest effect.

**Figure S5.**
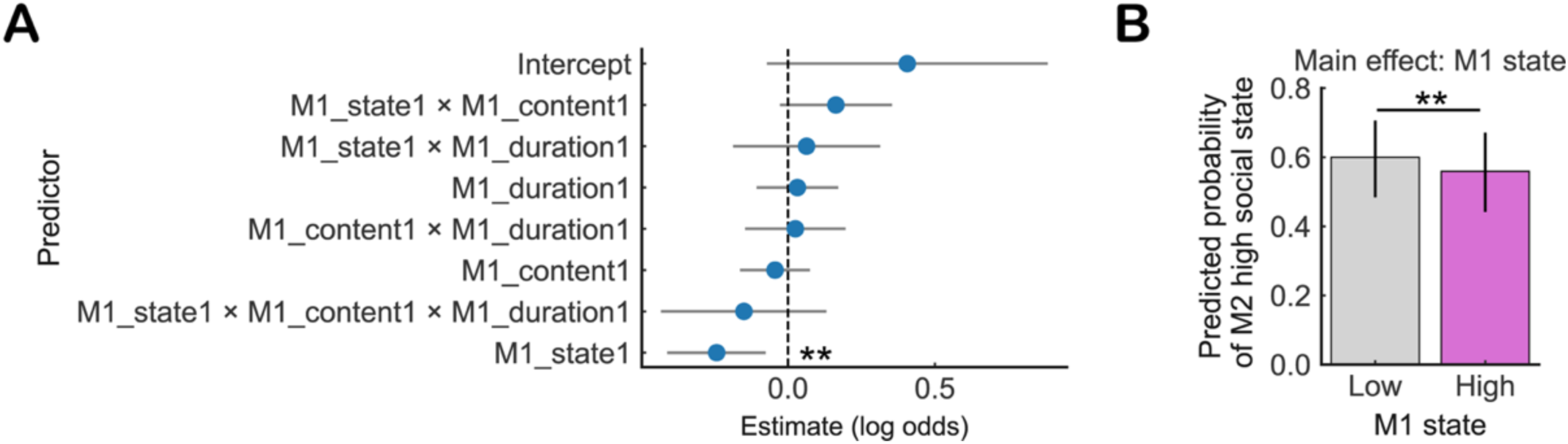
M1’s social state negatively predicts M2’s social engagement. Related to Fig. 1D. **(A)** Estimates of fixed effects from the generalized linear mixed-effects model (GLMM) predicting the occurrence of M2’s social state based on M1’s social gaze primitives and their compositions (Methods). Each point represents a fixed effect (log-odds estimate) with 95% confidence intervals. Asterisks indicate significance levels based on Type III Wald chi-square tests. The reference conditions for all predictors were defined as M1 state = Low, M1 content = Object, M1 duration = Short. **(B)** Predicted probabilities (back-transformed from the logit scale) of M2 high social state as a function of M1 content. Vertical lines on the bars indicate 95% confidence intervals. Horizontal lines with asterisks denote significant pairwise differences. Significance was tested using pairwise Wald z-tests on marginal mean differences (Tukey-adjusted), which account for covariance; thus, significance may occur even when 95% confidence intervals overlap. ** *P* < 0.01.

**Figure S6.**
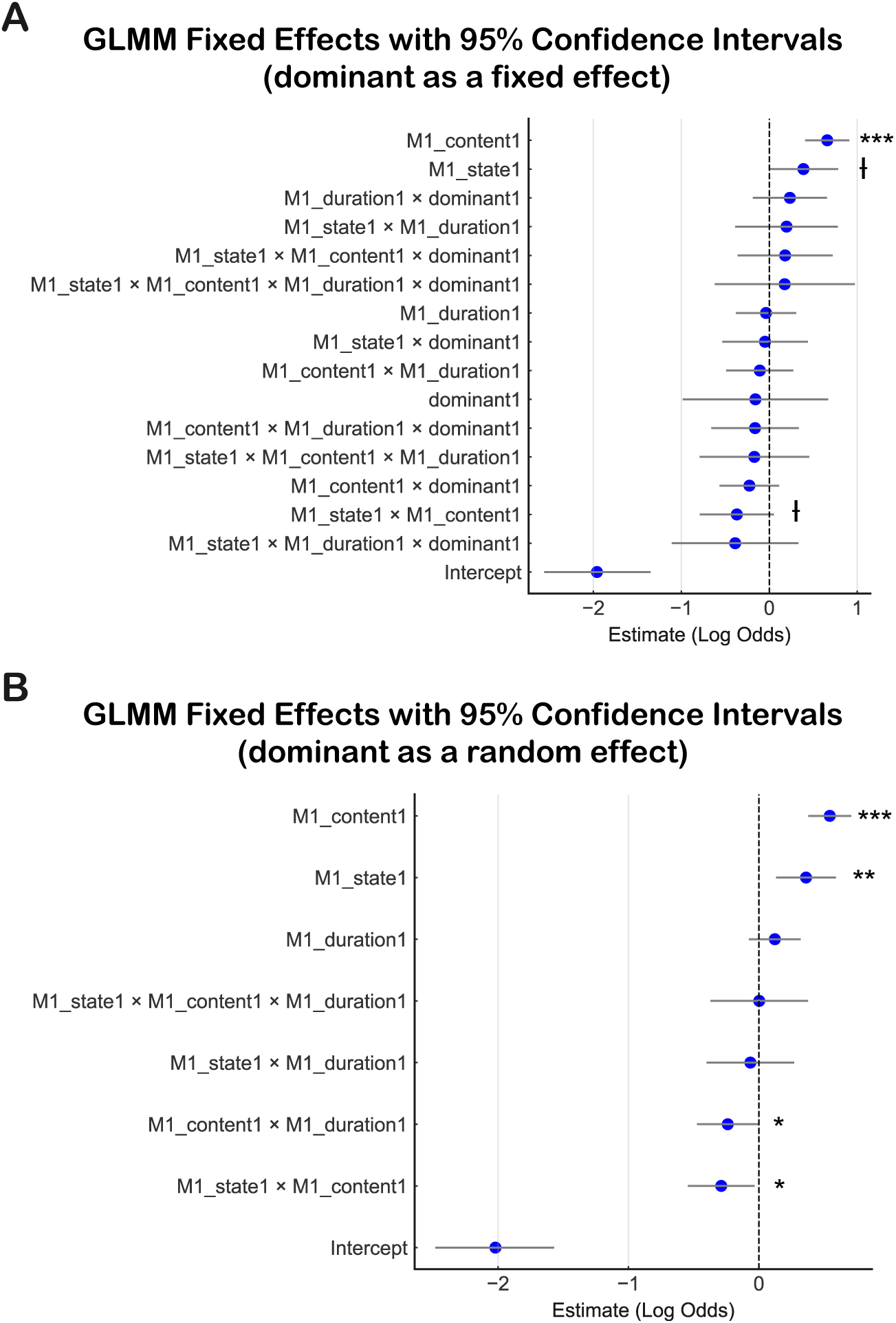
Partner dominance does not account for social gaze effects. Related to Fig. 1D. **(A)** Estimates of fixed effects from an extended generalized linear mixed-effects model (GLMM) predicting the occurrence of M2’s social gaze based on M1’s social gaze primitives (content, state, duration), partner dominance and their interactions. Points indicate fixed-effect log-odds estimates with 95% confidence intervals. Vertical dashed line indicates zero effect. The reference conditions for all predictors were defined as M1 state = Low, M1 content = Object, M1 duration = Short, and M1 dominance = 0. In this model, random intercepts were included for session/day, M1 identity, and M2 identity. **(B)** Fixed-effect estimates from a complementary GLMM in which dominance was modeled as a random intercept rather than a fixed effect. In this model, random intercepts were included for session/day, M1 identity, M2 identity, and M1 dominance. Points indicate fixed-effect log-odds estimates with 95% confidence intervals. Vertical dashed line indicates zero effect. Reference conditions were M1 state = Low, M1 content = Object, and M1 duration = Short. Significance was assessed using Type III Wald chi-square tests. Ɨ *P* < 0.1; * *P* < 0.05; ** *P* < 0.01; *** *P* < 0.001.

**Figure S7.**
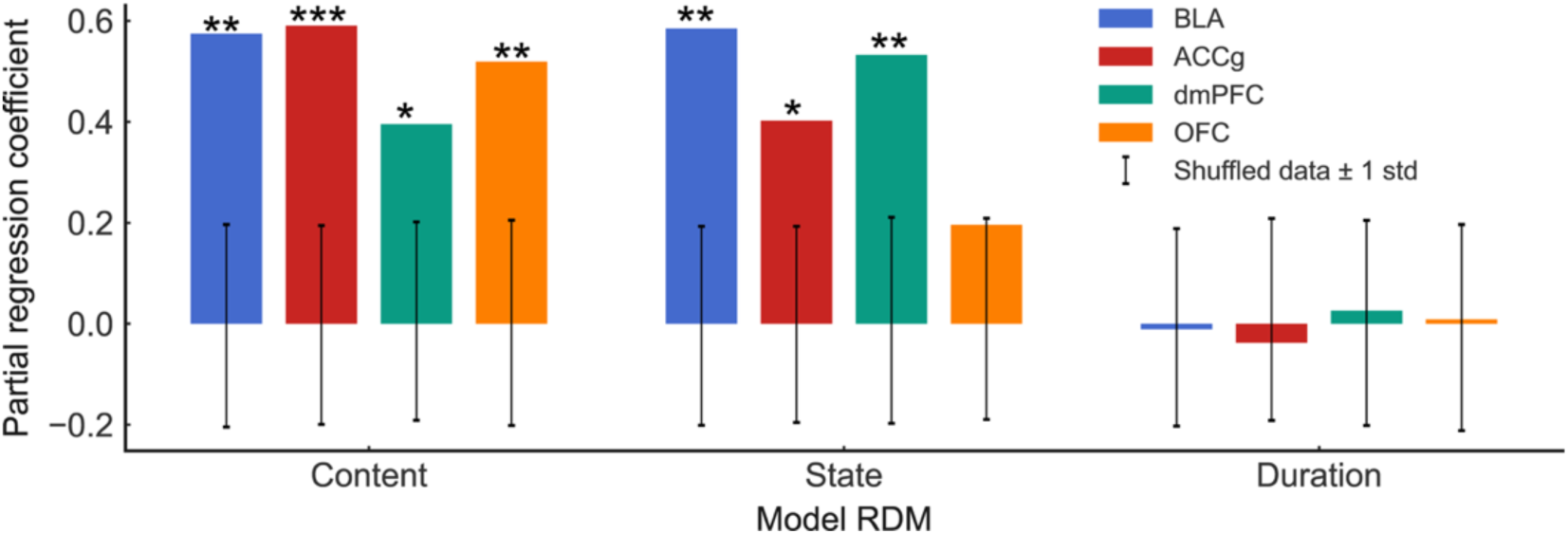
Encoding of the three social gaze primitives in the four neural populations based on a partial regression approach. Related to Fig. 2D. Shown are partial regression coefficients between brain RDM and social gaze primitives RDM. Bars show partial regression coefficients obtained from multiple regression of neural RDMs onto model RDMs, reflecting the unique contribution of each model after controlling for the others. * *P* < 0.05; ** *P* < 0.01; *** *P* < 0.001 with permutation test.

**Figure S8.**
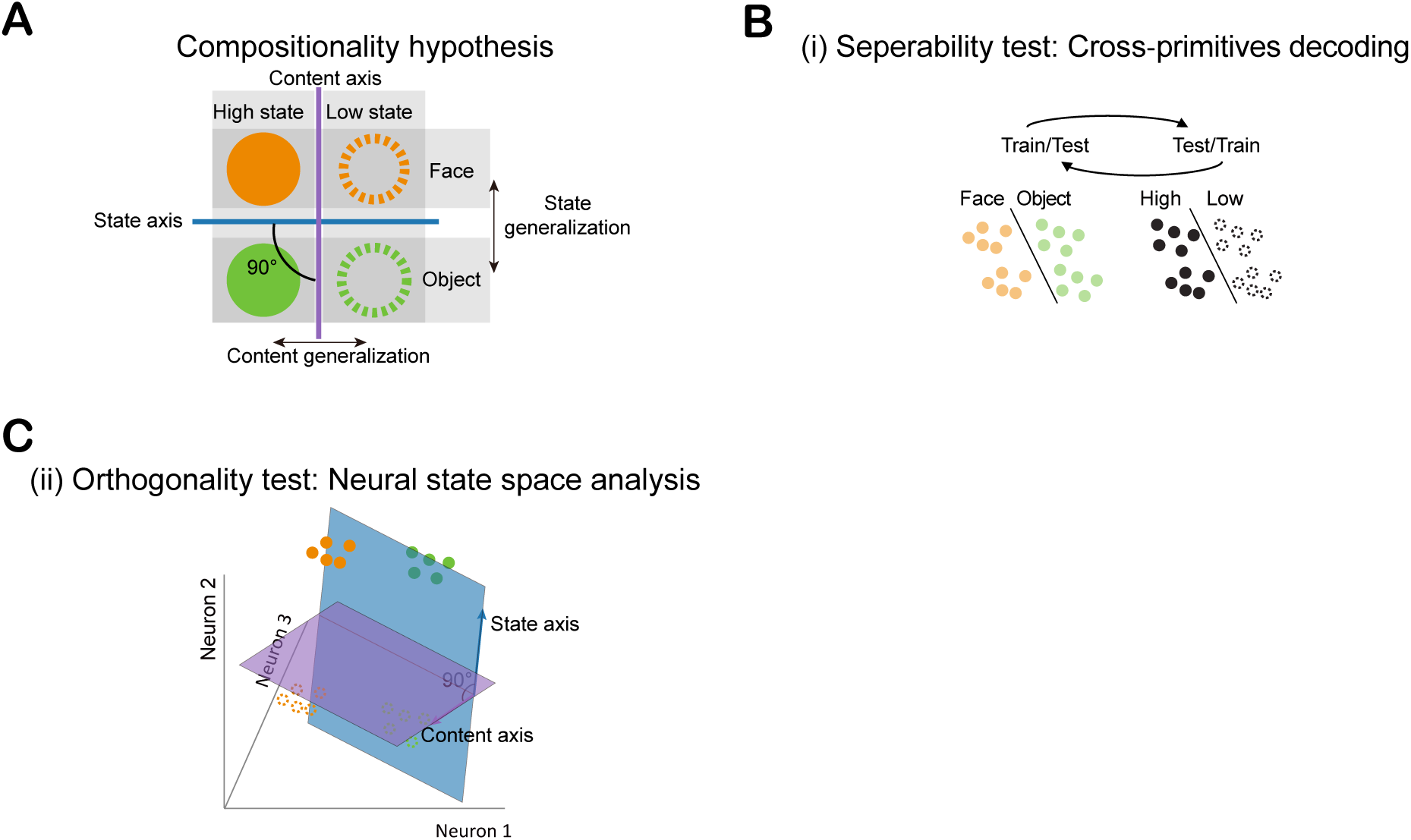
A hypothesis testing the compositionality of social gaze primitives at the neural population level with the three testable predictions. Related to Fig. 3. To evaluate whether neural populations represent social gaze in a compositional manner, we tested three predictions derived from the compositionality hypothesis: (i) social state and gaze content are represented distinctly; (ii) their neural encoding axes are orthogonal; and (iii) these representations generalize across conditions. Illustrated are theoretical schematics and population-level decoding analyses used to test these predictions. (A) Schematic of a compositional encoding hypothesis. Neural population responses to four conditions (content/state: face/high, object/high, face/low, object/low) are distributed along orthogonal axes in neural state space. Color (orange = face, green = object) and fill (solid = high state, open = low state) distinguish the two dimensions. (B) Cross-primitives decoding for separability test. A linear SVM classifier trained to decode one variable (e.g., content: face vs. object) using content-labeled gaze events was tested on trials labeled by the other variable (e.g., state: high vs. low), and vice versa. (C) Neural state space analysis for testing orthogonality. Content and state encoding axes were defined by condition-wise firing rate differences. An angle close to 90 between them indicates content and state along the orthogonal axis.

**Figure S9.**
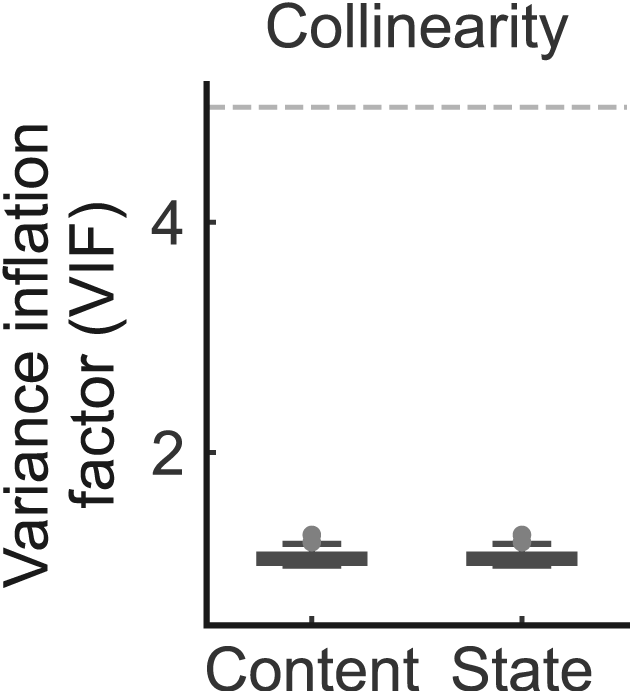
Independence of gaze content and social state predictors. Related to Fig. 4A. Variance inflation factor (VIF) distributions for the content and state predictors, computed at the single-neuron level. Both predictors show VIF values close to 1 and well below the conventional collinearity threshold (dashed line, VIF = 5), indicating negligible multicollinearity.

**Figure S10.**
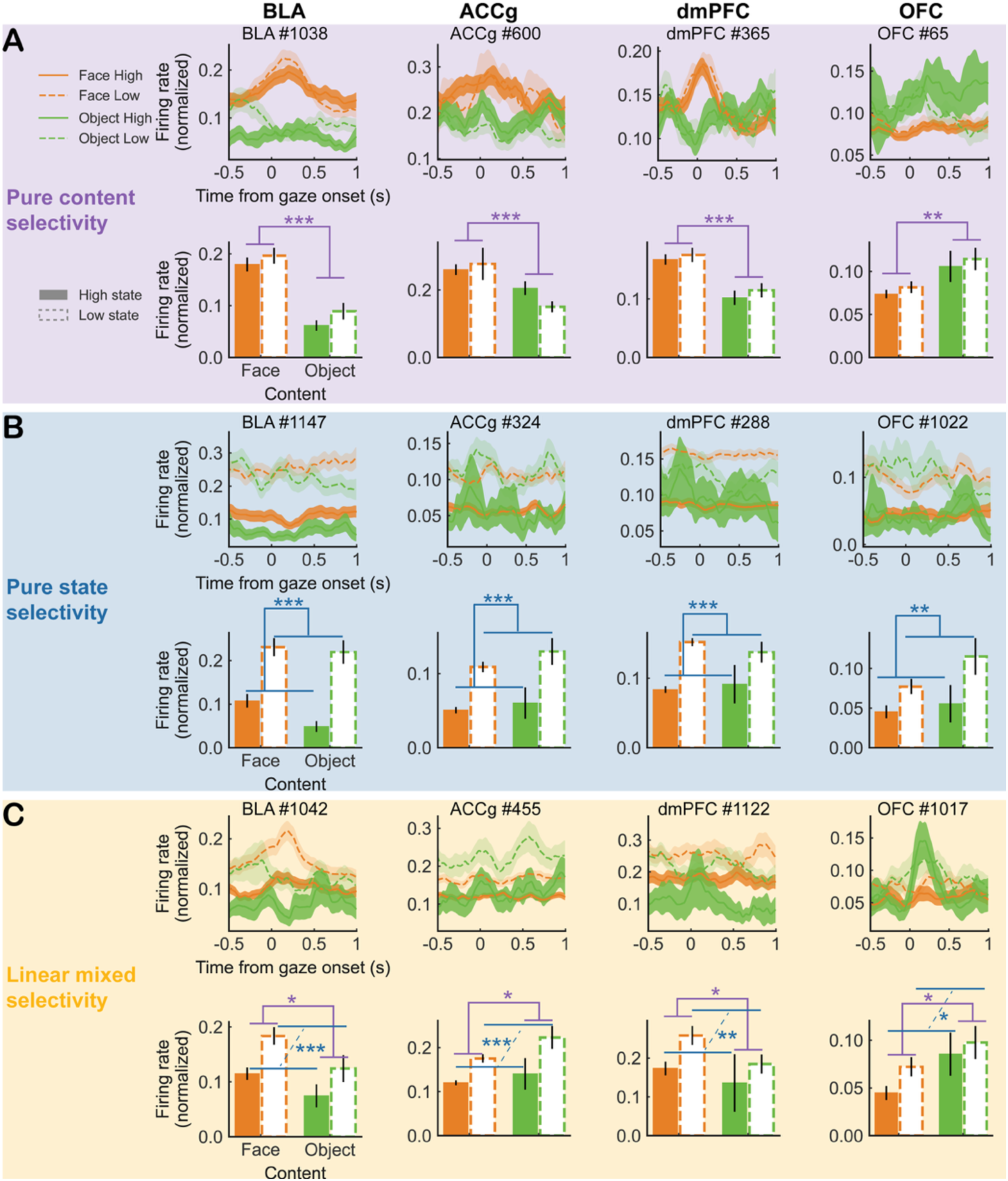
Example PSTHs for pure content-, pure state-, and linear mixed selectivity neurons in BLA, ACCg, dmPFC, and OFC. Related to Fig. 4B-C. Peri-stimulus time histograms (PSTHs) show normalized firing rates (min-max normalized to the [0, 1] range within each neuron) aligned to gaze onset (time 0) for example neurons classified as pure content-selective (A), pure state-selective (B), and linear mixed-selective (C). Each column shows representative neurons from a different brain region for each classification, respectively. For each representative neuron, its averaged firing rate during gaze events in each condition (face in high state, face in low state, object in high state, and object in low state) was plotted below its corresponding PSTH. Horizontal brackets indicate significant main effects of either gaze content (face vs. object, purple) or social state (high vs. low, blue) based on linear regressions in Fig. 4A. * *P* < 0.05; ** *P* < 0.01; *** *P* < 0.001. For the bar plots, error bars represent mean ± SEM across trials. For the PSTHs plots, shaded areas represent ±SEM across trials. Orange and green curves correspond to face and object gaze targets, respectively, with solid lines indicating high-state trials and dashed lines indicating low-state trials. Face High: solid orange; Face Low: dashed orange; Object High: solid green; Object Low: dashed green.

**Figure S11.**
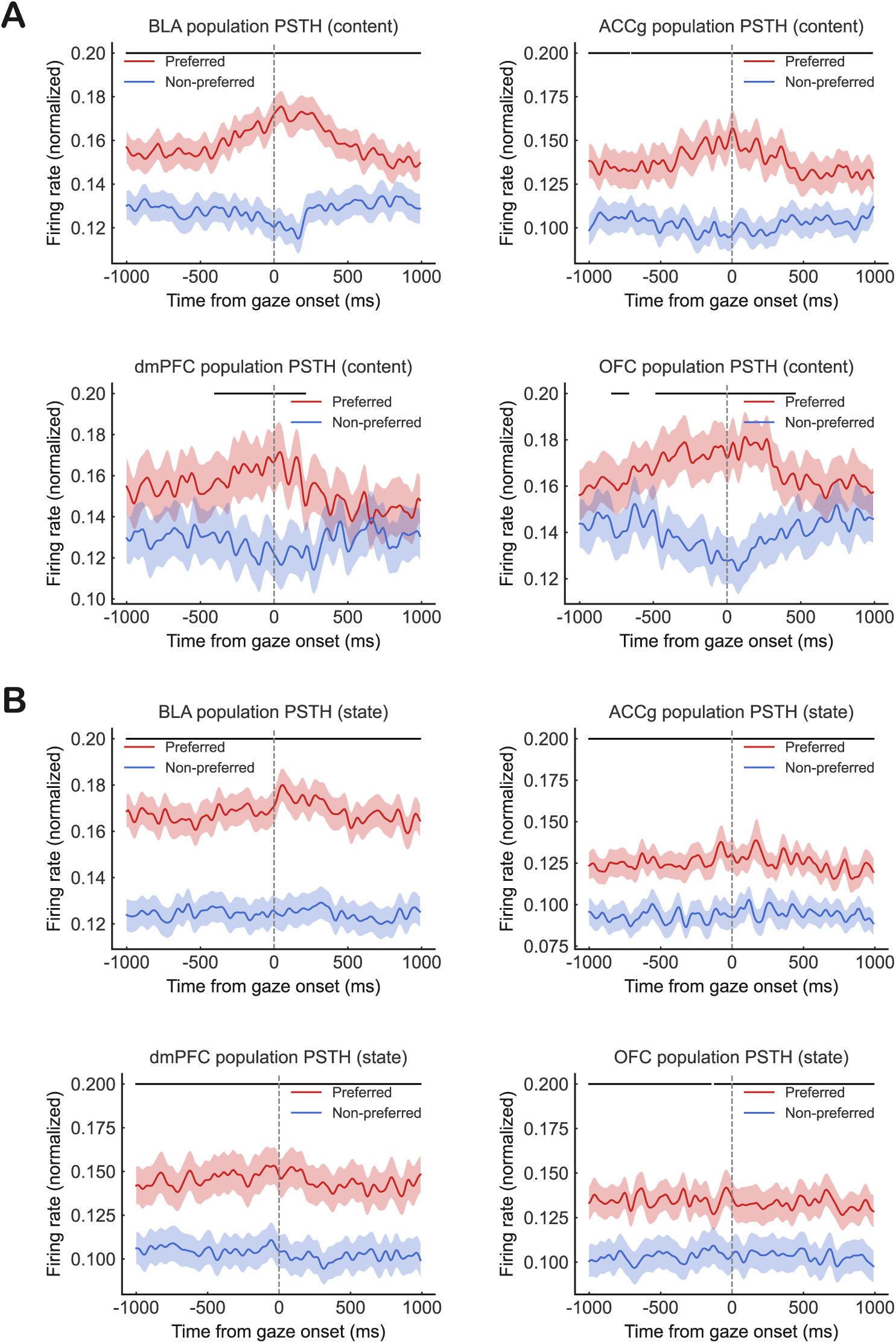
Population PSTHs recoded by preference for gaze content and social state. Related to Fig. 4B. **(A)** Population peristimulus time histograms (PSTHs) for neurons modulated by gaze content in BLA, ACCg, dmPFC, and OFC. **(B)** Same as **(A)**, but for neurons modulated by social state. PSTHs show normalized firing rates (min–max normalized to the [0, 1] range within each neuron) aligned to gaze onset (vertical dashed line at 0 ms). For each neuron, trials were split into independent training and test sets; the preferred condition (red) was defined on the training set and PSTHs were computed on held-out trials. Traces show the population mean ± s.e.m. across neurons. Black horizontal bars indicate significant temporally contiguous clusters in which preferred and non-preferred population responses differed (cluster-based permutation test, *P* < 0.05, minimum cluster duration 100 ms).

**Figure S12.**
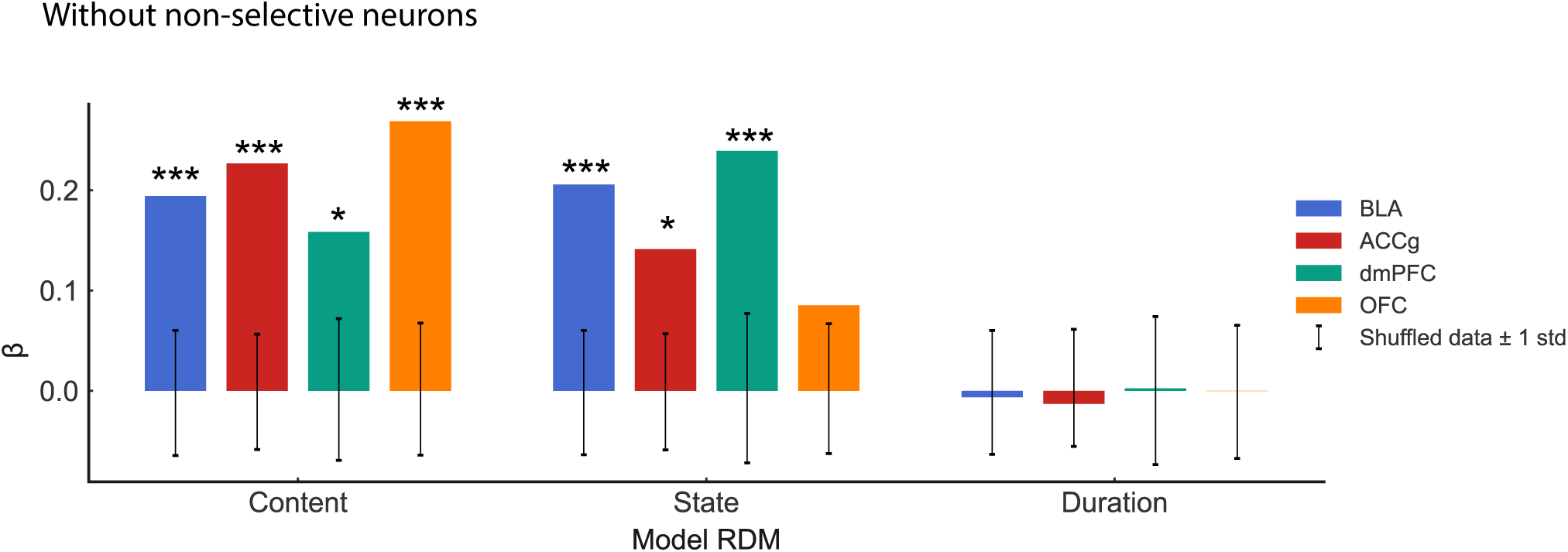
Encoding of the three social gaze primitives in the four neural populations (*without non-selective neurons*) based on the encoding GLM model. Related to Figs. 2D and 4B. Ɨ *P* < 0.1; * *P* < 0.05; ** *P* < 0.01; *** *P* < 0.001 with permutation test.

**Figure S13.**
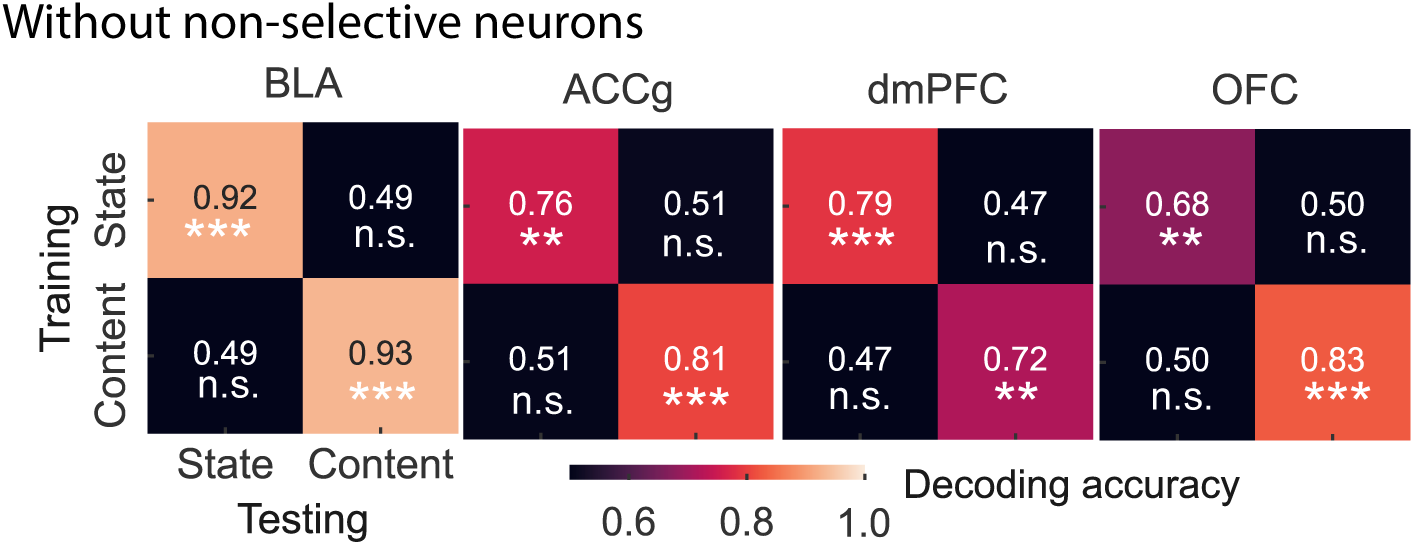
Decoding and cross-primitive decoding of content and state with four neural populations without non-selective neurons. Related to Figs. 3B and 4B. n.s., not significant; * P < 0.05; ** P < 0.01; *** P < 0.001 with permutation test.

**Figure S14.**
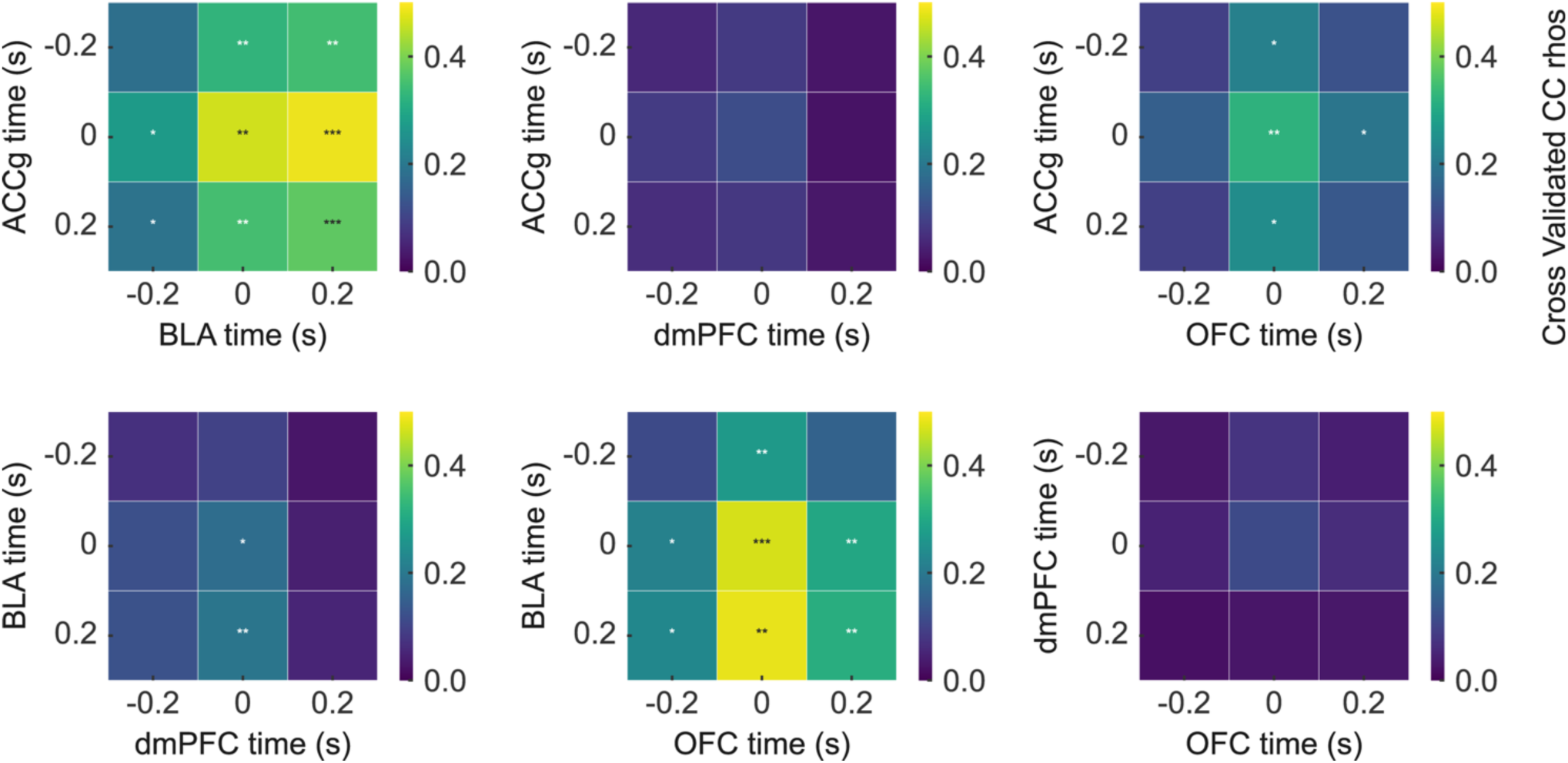
Averaged cross-temporal correlation matrices for content information. Related to Fig. 5A. * *P* < 0.05; ** *P* < 0.01; *** *P* < 0.001 with permutation test.

**Figure S15.**
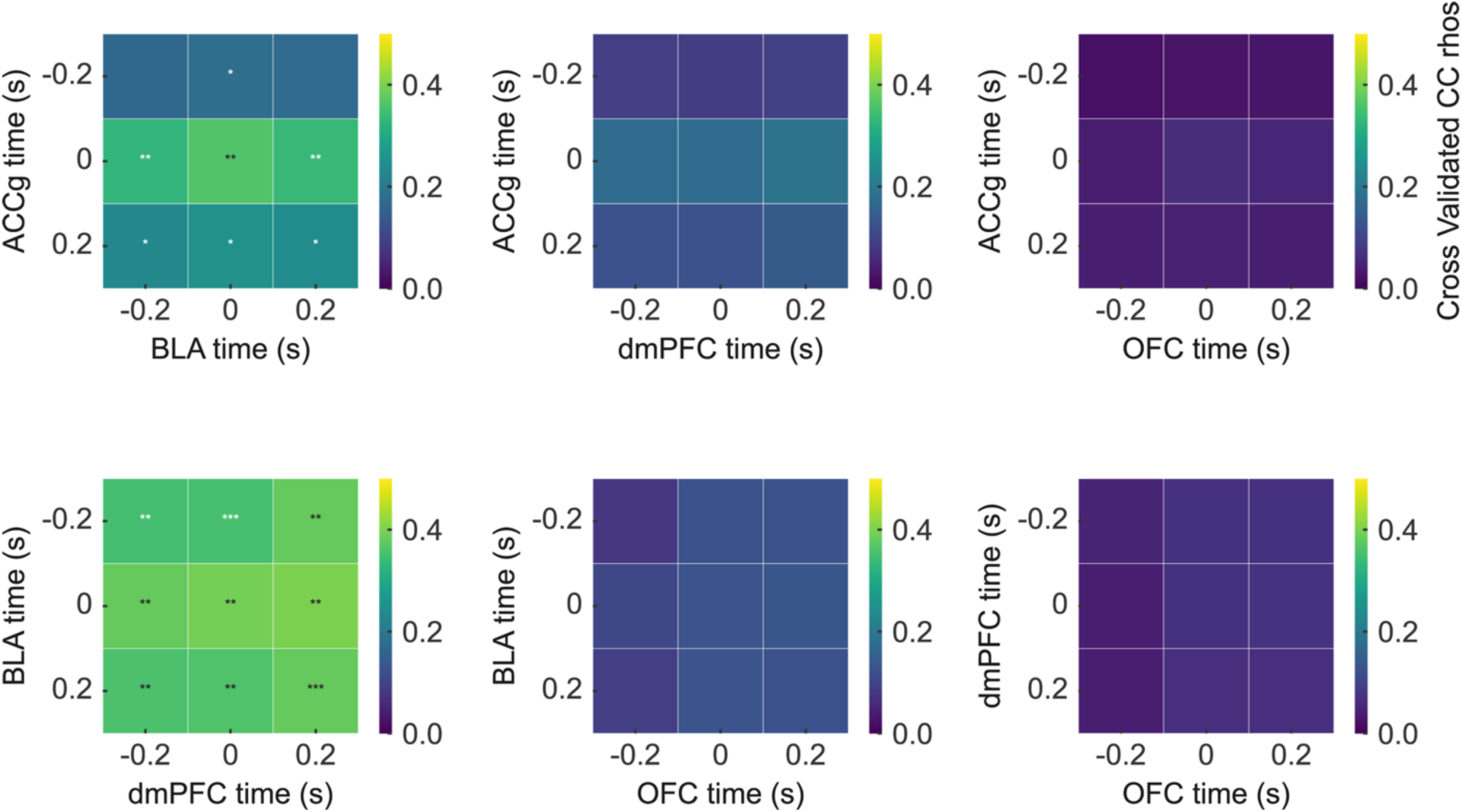
Averaged cross-temporal correlation matrices for state information. Related to Fig. 5A. * *P* < 0.05; ** *P* < 0.01; *** *P* < 0.001 with permutation test.

**Figure S16.**
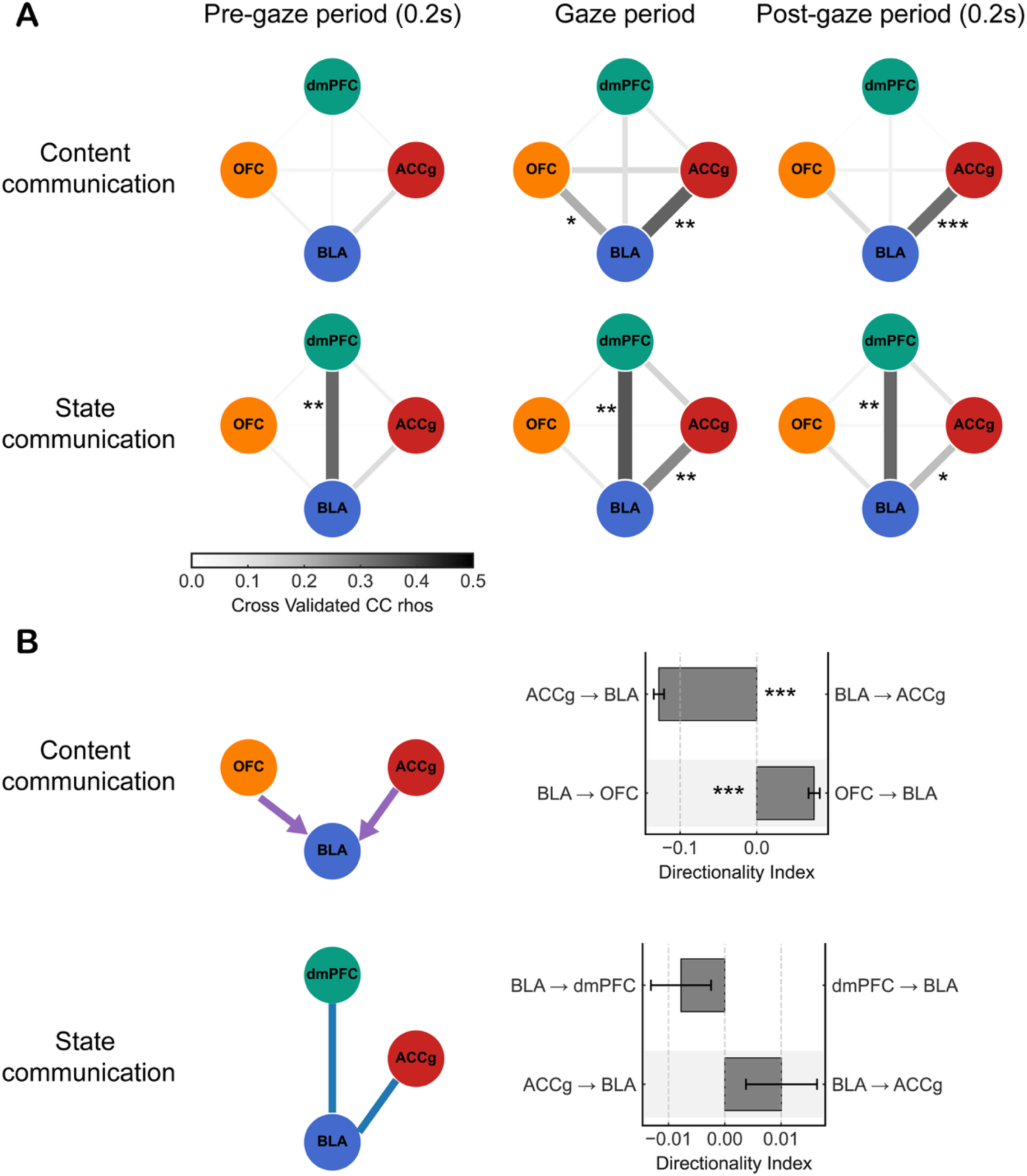
Functional connectivity of social gaze primitives when excluding linear mixed selectivity cells. Related Fig. 5. (A) Dynamic functional connectivity in the prefrontal-amygdala circuits for gaze content (upper panel) and social state (lower panel) information after removing linear mixed cells, respectively. Gaze period: when the eye gaze is on the face or the object. Pre-gaze period: 200msec before the eye gaze is on the face or the object. Post-gaze period: 200 msec after the eye gaze is on the face or the object. * *P* < 0.05; ** *P* < 0.01; *** *P* < 0.001with permutation test. (B) Directionality of the content (upper panel) and the state (lower panel) information after removing linear mixed cells for each pair of brain regions. Error bar: mean ± SEM. *** *P* < 0.001 with Wilcoxon signed-rank tests.

